# Single cell chronoatlas of regenerating mouse livers reveals early Kupffer cell proliferation

**DOI:** 10.1101/2021.06.09.447699

**Authors:** Daniel Sánchez-Taltavull, Tess Brodie, Joel Zindel, Noëlle Dommann, Bas G.J. Surewaard, Adrian Keogh, Nicolas Mélin, Isabel Büchi, Riccardo Tombolini, Paul Kubes, Daniel Candinas, Guido Beldi, Deborah Stroka

## Abstract

The liver is exemplar to study tissue regeneration due to its inherent ability of repair and regrowth. It replaces its lost or injured tissue by the proliferation, interaction and temporal coordination of multiple residential cell types. Until now we lacked a detailed description of the specific contributions of each cell type to the regenerative process, and therefore analyzed mouse livers 0, 3, 6, and 24 hours following two-thirds partial hepatectomy (PHx) by single cell RNA-sequencing (scRNA-seq) and mass cytometry. Our resulting genome wide temporal atlas contains the time dependent transcriptional changes in hepatocytes, endothelial cells, bone marrow-derived macrophages (BMDM) and Kupffer cells. In addition, it describes the cell specific contribution of mitogenic growth factors from biliary epithelial, endothelial and stellate cells as well as chemokines and cytokines from BMDM and granulocytes. And interestingly, Kupffer cells as opposed to hepatocytes emerged as the first cell to proliferate presenting a new dynamic in the liver following PHx. Here, we provide a robust data set at cellular resolution to uncover new elements and revisit current dogmas on the mechanisms underlying liver regeneration. To facilitate access to the data, we have launched the portal www.phxatlas.ch in which the scRNA-seq data can be visualized.

## Introduction

Maintenance of organ homeostasis and tissue renewal is critical for good health and longevity. Understanding of the mechanisms underlying tissue regeneration has the potential to improve the quality of human health and decrease deaths from end-stage organ failure^1^. Studies in the liver have been at the forefront of research in tissue regeneration because of its inherent ability of repair and regrowth. With the first rodent model of partial hepatectomy (PHx) published in 1931^2^, insight into the mechanisms of how the liver regenerates began to unfold^3^. Following the partial removal or damage to the liver, the remnant tissue initiates an immediate response to restore homeostatic levels of mass and function^4^ . Unlike other tissues which depend on stem cell replenishment, the liver restores its mass by the proliferation of its residential cell populations^5^. It has been widely accepted that the first cells to proliferate are the parenchymal hepatocytes which subsequently provide signals to the other liver cell populations^4^ . Technological developments such as the generation of knockout animals and RNA sequencing helped our understanding of this process by describing the contribution of many signaling molecules, cytokines and growth factors ^6, 7^. However, these factors are produced from various cellular compartments and bulk sequencing techniques are unable to define the source of the mRNA and therefore the exact contribution of each cell type remains unclear. Now, with the aid of high throughput single cell technologies, single cell RNA sequencing (scRNA-seq) and Cytometry by Time-of-Flight (CyTOF), we are able to provide further insight into the mechanism of regeneration at a cellular resolution. Here we report our data set on the dynamics of liver regeneration by mapping the cell specific transcriptional responses over time and uncover an unexpected insight.

## Results

### scRNA-seq and mass cytometry of the time dependent regulation in single cell populations after PHx

To study the transcriptional events in individual cells during liver regeneration, two-thirds of the liver was surgically removed by PHx. The regenerating lobes were digested 0, 3, 6 and 24-hours following surgery and single cell suspensions of parenchymal and non-parenchymal cell fractions were sequenced using 10x Genomics and Illumina sequencing (Fig. 1a). Following the scRNA-seq data preprocessing, the data set comprised the transcriptomic profile of 11’896 cells over four time points (Extended Data Table 1).

**Fig. 1.**
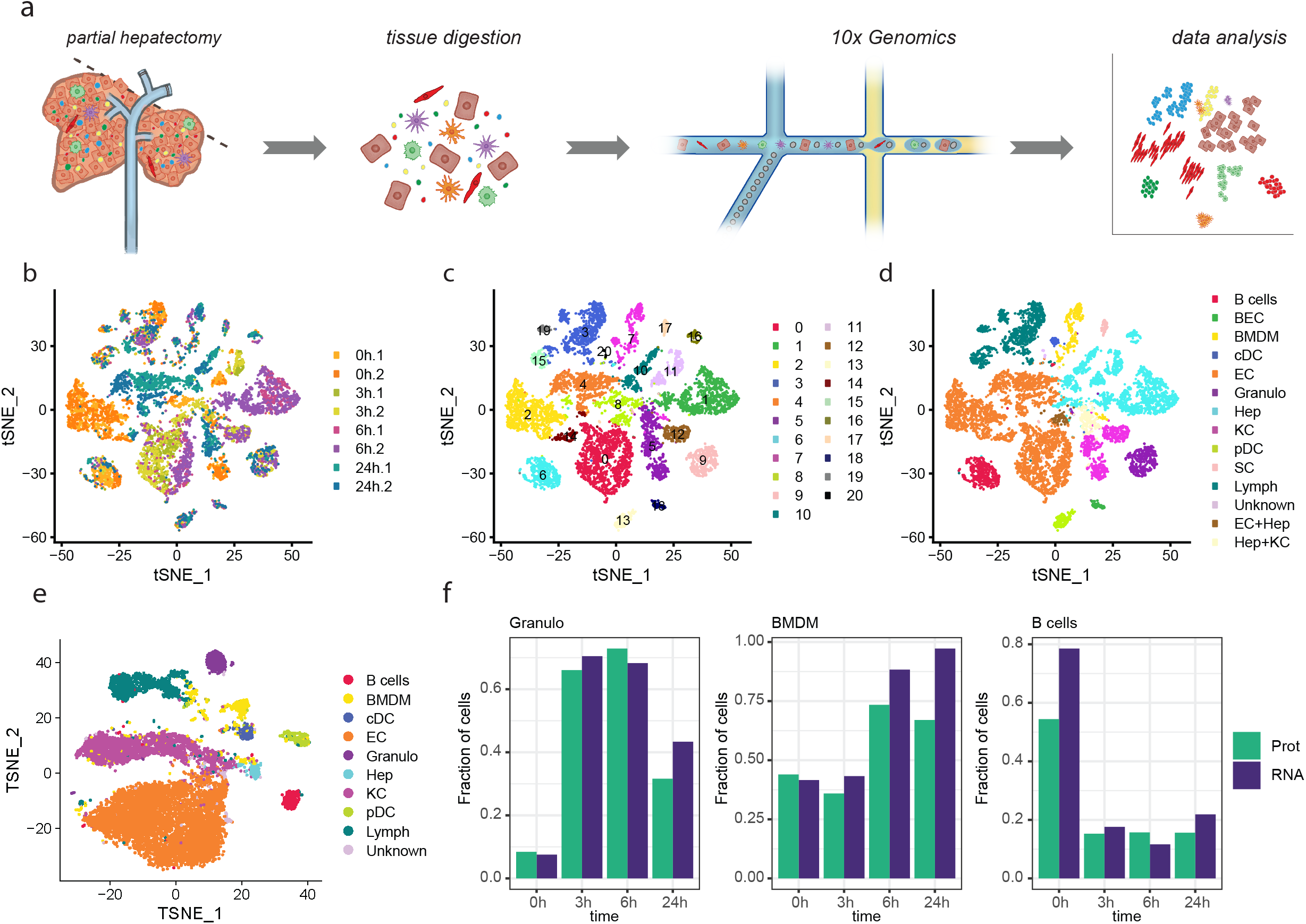
Single cell data description. a) Graphical display of experimental design. Liver cells were isolated with an in vivo collagenase digestion at time points 0, 3, 6 and 24h post PHx in duplicate, followed by 10x genomics library preparation and Illumina sequencing. b) t-SNE visualization of scRNA-seq colored by sample, c) by unsupervised cluster, d) by cell type. e) t-SNE visualization of CyTOF colored by cell type f) Normalized fraction of cells found in scRNA-seq (purple) and CyTOF (green).

The t-Distributed Stochastic Neighbor Embedding (t-SNE) visualization of the filtered scRNA-seq data showed the appearance of time dependent clusters an observation that was highly consistent between replicates (Fig. 1b). Unsupervised clustering identified 21 clusters (Fig. 1c). Using marker expression these 21 clusters were designated as either hepatocytes, endothelial cells, biliary epithelial cells, stellate cells, and immune cells including T, NK and NKT cells, B cells, granulocytes, classic and plasmacytoid dendritic cells and bone marrow-derived macrophages (BMDM) and Kupffer cells (Fig. 1d and Extended Data Fig. 1 & 2). Doublets were identified by marker expression of two or more cell types on a single cell and these cells were excluded from further analyses (Extended Data Fig. 3).

We next followed changes in the populations with a 33 parameter CyTOF panel designed mainly for the immune cell populations. By CyTOF, we detected all cell populations identified by scRNA-seq, with the exception of stellate and biliary epithelial cells (Fig. 1e, Extended Data Fig. 4). The CyTOF data confirmed a time dependent regulation in the endothelial cells and Kupffer cells (Fig. 1e, Extended Data Fig. 5). By comparing the scRNA-seq and CyTOF values of lineage marker expression, we observed a strong correlation between average mRNA and protein expression among the cell types, with a positive correlation for most genes (Extended Data Fig. 6a, b). In addition, we observed a change in the frequency of some cell populations that was consistent between the mRNA and protein data sets. For example, there was a time dependent increase in the number of granulocytes and BMDM, while B cell numbers decreased (Fig. 1f, Extended Data Fig. 7, 8). There were no significant differences in cell numbers of the other populations over time (Extended Data Fig. 7, 8).

### Cell specific transcriptional response to PHx

We next focused on cell specific transcriptional changes and performed a hierarchical clustering with the 50 most significant markers of each unsupervised cluster. The data grouped first by cell type and then in some populations by time (Extended Data Fig. 9). To study changes over time, we analyzed each cell population separately, and facilitate access to the gene expression profiles for each population through the portal www.phxatlas.ch to visualize the data.

Macrophage activation is an early and initiating event required for the liver to regenerate^8^. Therefore, we began by studying the profiles and potential cell-cell interactions of two residential macrophage populations which separated into pro-inflammatory BMDM and yolk sac–derived, immunoregulatory Kupffer cells ^9–11^. These two populations are similar to those observed in humans with cell markers consistent between the human and mouse liver ^12^. Both populations showed a time dependent transcriptional regulation following PHx (Fig. 1d, e, Fig. 2a).

**Fig. 2.**
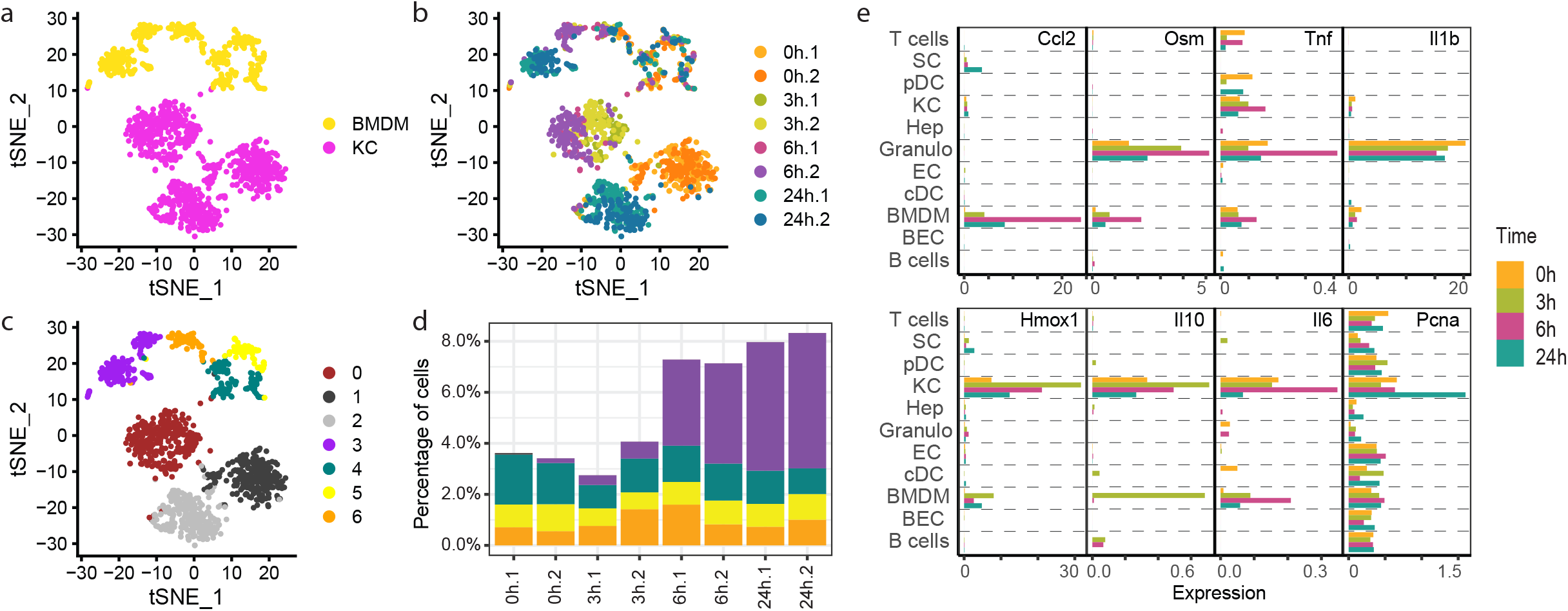
Cell specific transcriptional responses to PHx. A) t-SNE visualization of the BMDM and KC colored by cell type, b) by sample, c) by unsupervised cluster. d) Stack plot showing the percentage of each unsupervised cluster of BMDM present in every sample. e) barplot of the gene expression of Ccl2, Osm, Tnf, Il1b, Hmox1, Il10, Il6 and Pcna in the different cell types.

The BMDMs separated into five clusters (Extended Data Fig. 10a, b, c) with new clusters emerging at 6 and 24 hours (Fig. 2b, c, d). The mRNA dynamic was supported by CyTOF data (Extended Data Fig. 11). The newly emerged clusters coincided with their increased abundancy (Fig. 1f) and their transcriptomic profile identified pathways associated with acute inflammatory responses and protein processing (Extended Data Fig. 12). Genes increased included many chemokines, such as *Ccl2* (Fig. 2e), *Ccl6*, *Ccl7* and *Ccl9* (Extended Data Fig. 13a). Plausible communication mediating migration and infiltration was found between the recruited BMDM via *Ccl2* and *Ccl7* and their receptor *Ccr2* (Extended Data Fig. 13a, Extended Data Table 2). In addition, communication between *Ccr1* found on macrophages and granulocytes^6^ and *Ccl9* may contribute to the trafficking of these two cell types to the liver (Extended Data Fig. 13a, Extended Data Table 2). Inflammatory cytokines, such as oncostatin M (*Osm*) and tumor necrosis factor (*Tnf*) were increased at 6 hours in BMDM (Fig. 2e), interestingly, however, the highest expression of *Osm* and *Tnf* was found in the granulocytes (Fig. 2e). Granulocytes were also the main producer of *Il1b* (Fig. 2e).

Within the Kupffer cells, three clusters emerged with changes in their transcriptional profile differing quite remarkably from the BMDM (Fig. 2a, b, c, Extended Data Fig. 14a, b, c). One cluster contained the cells associated with the 3 and 6-hour timepoints. Within this cluster, pathways such as responses to external stimuli and shear stress were prominent (Extended Data. Fig. 15), in which anti-oxidants heme oxygenase (*Hmox1*; Fig. 2e) and metallothioneins (*Mt1* and *Mt2*; Extended Data Fig. 13b) were among the highest differentially expressed genes. Even with a significant increase of *Ccl6* and *Ccl9* (Extended Data Fig. 13a), pathways associated with acute inflammatory response and cytokine-mediated signaling pathways were largely unchanged over time (Extended Data Fig. 15). The highest levels of the Th2 cytokine *IL10* was found in Kupffer cells (Fig. 2e). Although *IL6* signaling is essential for regeneration by inducing an acute phase response^7, 13^, a low level of *IL6* expression was found in Kupffer cells and BMDM (Fig. 2e). The most striking change in Kupffer cells occurred in the cluster associated with the 24-hour time point in which proliferation pathways prominently appeared (Extended Data Fig. 15). In this cluster, there was an increased expression of DNA replication genes such as *Pcna* (Fig. 2e), *Cdt1, Mcm5* and *E2f1* (Extended Data Fig. 16).

To compare the novel finding of a transcriptional increase of DNA replication genes in the Kupffer cells at 24 hours, we next checked the other cell populations. Transcriptional changes in biliary epithelial, stellate cells, granulocytes, B cells and dendritic cells were not enough to separate these cells further into time-dependent clusters (Extended Data Fig. 17-22). For the biliary epithelial and stellate cells, this may be due to the low number of cells sequenced and therefore their potential contribution to regeneration is missed. In the lymphocyte population, one cluster separated (Extended Data Fig. 23) whereas, in the hepatocyte and endothelial cells several time dependent clusters emerged. Six hepatocyte clusters were identified, of which five correlated with time (Extended Data Fig. 24a, b, c). A pathway analysis of the clusters containing cells at the 3, 6 and 24-hour time points indicated transcriptional changes mainly related to changes in metabolic pathways and no pathways related to genes involving DNA replication were observed. (Extended Data Fig. 25). Six different endothelial cell clusters were identified with four of the clusters showing a strong correlation with time (Extended Data Fig. 26a, b, c). The transcriptomic profile of endothelial cells at 3 hours indicated a response in pathways related to hormones and cytokine production and blood vessel morphogenesis. The key pathways regulated at 6 and 24 hours included those involved in inflammatory responses, binding and uptake of ligands by scavenger receptors and regulation of cell adhesion (Extended Data Fig. 27). Likewise, no pathways related to genes involving DNA replication were observed. Of note, the endothelial cells expressed the highest mRNA levels of the inflammatory cytokine *Il1a* (Extended Data Fig. 13c).

### Validation of Kupffer cell proliferation

The observation that pathways associated with proliferation were the most regulated pathways in Kupffer cells at 24 hours suggested that they were dividing, thus challenging the current dogma that hepatocytes are the first cells to proliferate following PHx^14, 15^. To experimentally confirm this, we first compared DNA replication genes across the entire data set and observed that only Kupffer cells they were increased at 24 hours (Kupffer cells and hepatocytes, Fig. 3a, all other cell types, Extended Data Fig. 28). Kupffer cells were then categorized based on cell cycle marker expression and most were assigned to the S and G2M phase at 24 hours (Fig. 3b, Extended Data Fig. 29). Next, we questioned if the thymidine analogue, 5-ethynyl-2-deoxyuridin (EdU) was incorporated into Kupffer cells between 24-26 hours post PHx. There was a significant increase in EdU positive Kupffer cells suggesting they were in DNA synthesis/S-phase (Fig. 3c, Extended Data Fig. 30). Finally, we used intravital confocal spinning disc microscopy with a Clec4F driven reporter system to identify the cytoplasm (CS green) and nuclei (dtTomato) of Kupffer cells to visualize Kupffer cell proliferation. Time-lapse imaging demonstrated that Kupffer cells indeed undergo cell division at 24 hours (Fig. 3d, Supplementary Video 1).

**Fig. 3.**
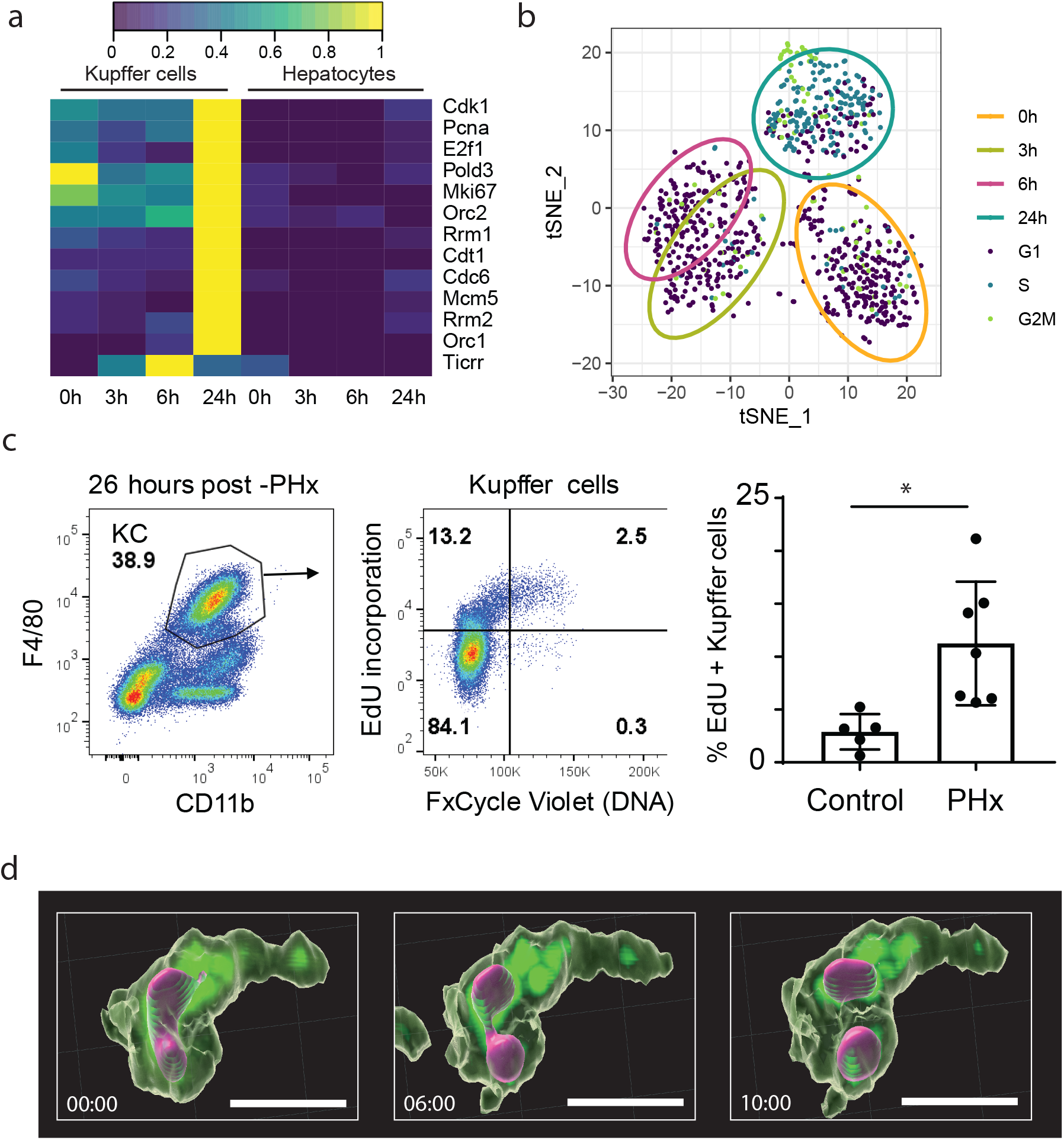
Validation of KC proliferation. a) Heatmap of the average mRNA expression of DNA replication genes in Kupffer cells and hepatocytes. b) t-SNE visualization of cell cycle phase assigned based on the cell cycle score (see methods) of each KC. The ellipses show the interval of confidence of each time point. c) Murine non-parenchymal liver cells label with EdU 24-26 hours post PHx. Gated F4/80hi, CD11b+ Kupffer cells were analyzed for EdU incorporation and DNA content. d) Spinning disk confocal in vivo microscopy of the regenerating right liver lobe 24h post PHx. Red tdTomato nuclear staining. The mouse expressed a Cre under the Clec4F promotor and was crossed to a stop-floxed GFP-reporter.

As growth factors are primary mitogens for cell proliferation^16–18^, we checked for expression and potential interaction of growth factor receptor signaling in Kupffer cells. We observed that endothelial, biliary epithelial and stellate cells were the main source of most of the growth factors, such as *Tgfa, Hbegf, Hgf, Pdgfs, Fgf1* and *Vegf*, with the endothelial cells the most transcriptionally reactive to PHx (Extended Data Fig. 31). Of the growth factor receptors, *Met, Fgfr1, Pdgfra, Tgfbr1, Flt1*, only *Egfr* expression was increased in Kupffer cells at 6 and 24 hours (Extended Data Fig. 31). As *Egf* mRNA levels were relatively low in all liver populations (Extended Data Fig. 31a), most likely due to its extrahepatic production^19^, it could not be assigned as the primary mitogen for Kupffer cell proliferation from this data set.

## Discussion

With the advancements in single cell technologies a more in depth description of liver cell populations under homeostatic and diseased conditions has been possible^12, 20, 21–23^. Here we are able to provide a deeper insight to the complex process of liver regeneration by documenting in a chronoatlas the transcriptional contribution of individual cell populations of the liver over time. One of our most interesting observation was that Kupffer cells are the first cell type to proliferate following PHx. This observation is contrary to current knowledge suggesting Kupffer cell division follows hepatocyte expansion^15^. It does however support that Kupffer cells are maintained by local proliferation at steady state^24^ and does not dispute the possibility that Kupffer cells can be replenished by peripheral BMDM as they are not depleted in our model of PHx^25, 26^. The importance of Kupffer cells in liver regeneration is well documented, for example, in clondronate-mediated depletion of Kupffer cells, liver regeneration is delayed and there is an increase in liver damage following PHx^27, 28^. However, these depletion studies have mainly been used to denote the importance of Kupffer cells due to the coinciding loss of cytokine availability. Early KC proliferation, may indicate that they are providing additional vital functions during regeneration, for example, optimal numbers may need to be maintained in order to cope with the upcoming expansion of parenchymal cells. Kupffer cells are located in the hepatic sinusoids, not in direct contact with hepatocytes. Their close proximity and cross talk with endothelial cells support that Kupffer cells may have a role in helping to maintain vascular integrity during a process of dynamic changes in blood flow and pressure^29, 30^. Further studies are underway to understand the importance of early proliferation, for example, to address the question if Kupffer cell proliferation is a prerequisite for hepatocyte division.

## Acknowledgements

We would like to acknowledge the Next Generation Sequencing Platform at the University of Bern. We would like to thank Helena Crowell for her CATALYST package in R that was used to analyze the CyTOF data and for her support with questions on the analysis. We gratefully acknowledge Dr. Anna Alemany and Aram Ortega Adzerias for fruitful discussions.

## Methods

### Partial hepatectomy

The mice used in this study were 12-week male C57BL/6JRj (Janvier Labs, France). Animals were kept under SPF conditions in a temperature-controlled room with a 12-h dark/light cycle. Littermates were randomly assigned to control or partial hepatectomy groups. Mice were anesthetized by isoflurane inhalation during an operation time of 15–20 min. Briefly, following a laparotomy, vicryl 4.0 suture (VCP496, Ethicon) was used to ligate and excise the median and left liver lobes. The peritoneal cavity was washed with saline solution and the abdomen was sutured closed. For analgesia, Temgesic (0.05 mg/kg) was injected i.p. prior to surgery. Experiments were done with Institutional Animal Care and Use Committee approval and in strict accord with good animal practice as defined by the Office of Laboratory Animal Welfare.

### Cell isolation

Animals were anesthetized (fentanyl 50ug/kg, midazolam 5mg/kg, medetomidine 500ug/kg, i.p.), immobilized in a supine position and the liver and portal vein exposed. The portal vein was cannulated with a 22G catheter and perfusion at 4ml/min with the buffers allowed to run to waste through an incision in the inferior vena cava. The liver was perfused with 10 ml of HBSS (Mg2+, Ca2+ free,10mM HEPES, pH7) followed by 25ml of HBSS containing EDTA (10mM HEPES pH7, 5mM EDTA). The EDTA was removed from the liver by perfusion with 10 ml of HBSS followed by digestion with 25 ml of HBSS containing collagenase (10mM HEPES, 1mM CaCl2, 0.5mg/ml collagenase IV, 0.01mg/ml collagenase 1A (Sigma)). The liver was excised and the cells released by cutting the capsule and gently agitating the digested liver in stop buffer (HBSS, 10mM HEPES pH7, 5mM EDTA, 10mM citrate, 1% FBS) and passed through a 70μm filter. The cell suspension was centrifuged at 30g min to pellet most of the hepatocyte fraction, the supernatant was collected and remnant cells pelleted at 250xG. This cell pellet was washed once in stop buffer then re-suspended in 20% isotonic Percoll and overlaid on a layer of 80% isotonic Percoll and centrifuged at 500G for 10 minutes. The cells at the interface of the two Percoll layers were collected and washed in PBS, re-suspended in PBS and counted with a cell counter (Bio-Rad, TC20). Hepatocytes from the initial 30G spin were then added to these cleaned cells to give a final hepatocytes concentration of approximately 10% of the total cell number.

### Library preparation

scRNA-seq libraries were prepared from 5000 cells from sham control, 3 hours, 6 hours and 24 hours after PH in duplicate using the Chromium Single Cell 3’ Library & Gel Bead Kit v3 (10xGenomics). Libraries were prepared according to the manufacturers protocol.

### Sequencing

Sequencing was performed on a NovaSeq 6000 S2 flow cell. Read 1 consisted of 26 cycles (10XGenomics barcode plus UMI) followed by a single Illumina i7 index read of 8 cycles and read 2 of 91 cycles to determine transcript-specific sequence information.

### Read alignment

The function cellranger count from Cell Ranger^31^ was used to transform the fastq files with the parameter *expect-cells* set to 5000. The reference genome was the mm10 available at Illumina Cell Ranger webpage. Next, we used the function *cellranger mat2csv* to generate the UMI matrix.

### Data pre-processing

First, cells expressing less than 200 genes were excluded. Second, dead cells identified as cells with more than 15% reads coming from mitochondrial genes (see Extended Data Table 3) were excluded. Third, erythrocytes identified as cells with more than 25% of reads coming from globin genes (see Extended Data Table 3) were removed. Fourth, genes not expressing at least 2 UMIs in at least 2 cells were excluded from the analysis.

After data cleaning, Ensembl IDs were transformed into mgi symbol with the R package biomaRt. The resulting filtered UMI matrices were transformed into Seurat objects with the function *CreateSeuratObject* with the UMI matrix as *counts, min.cells=1, min.features=1*. The 8 seurat objects were merged with the function merge.

The data was normalized, the highly variable genes were identified and scaled with the function *SCTransform* with parameters *vars.to.regress = “percent.mt”*, considering all the cells.

### Dimensionality reduction

We performed a Principal Component Analysis (PCA) of the Seurat object with the R function *RunPCA for all the cells, and for each gated subpopulation*. We performed a tSNE dimensionality reduction with the R functions FindNeighbors and *RunTSNE* with parameters *dims=1:30 (All cells), dims=1:8 (B cells), dims=1:10 (BEC), dims=1:20 (BMDM), dims=1:8 (cDC), dims=1:20 (EC), dims=1:12 (Granulo), dims=1:20 (Hep), dims=1:30 (KC), dims=1:10 (pDC), dims=1:10 (SC), dims=1:10 (T cells), dims=1:30 (macrophages), dims=1:10 (cluster 5), dims=1:20 (cluster 7), dims=1:12 (cluster 8).* Confidence ellipses were performed with the *stat_ellipse() from the R package ggplot2.*

### Unsupervised Clustering

We identified the unsupervised clustering with the R function *FindClusters* varying the parameter *resolution. resolution=0.35 (All cells), resolution= 0.15 (B cells), resolution= 0.15 (BEC), resolution= 0.2 (BMDM), resolution= 0.15 (cDC), resolution= 0.15 (EC), resolution= 0.2 (Granulo), resolution= 0.15 (Hep), resolution= 0.15 (KC), resolution= 0.1 (pDC), resolution= 0.2 (SC), resolution= 0.2 (T cells), resolution=0.5 (macrophages), resolution=0.5 (cluster 5), resolution=0.5 (cluster 7), resolution=0.5 (cluster 8).*

For all cells and for each individual cell type, the effect of the resolution was studied varying the parameters (Extended Data Fig. 32a,b, and Fig. 10d,e, Fig. 14d,e, Fig. 17-23d,e, Fig. 24d,e and Fig. 26d,e).

### Cell type identification and doublet removal

To study how the clustering grouped the cells, the entire data set was clustered with increasing resolution which led to the appearance of the identified cell types, and the time dependent regulation, resulting in the 21-cluster classification (Extended Data Fig. 32a, b). Increasing the resolution further led to the emergence of additional subtypes of endothelial cells, hepatocytes, BMDM, Kupffer cells, and T cells; however, still did not lead to a separation in B cells, classic or plasma dendritic cells, biliary epithelial, granulocytes, or stellate cell clusters (Extended Data Fig. 32a, b).

Cell types were identified based on marker expression (see Extended Data Table 4). Each unsupervised cluster identified with resolution=0.35 was assigned to a unique cell type, except for Clusters 5, 7 and 8. Cluster 5, containing KC and Hep, was reanalysed and a new unsupervised clustering was done to separate KC and doublets KC+Hep. Cluster 7, containing BMDM and cDC, was reanalysed and a new unsupervised clustering was done to separate BMDM and cDC. Cluster 8, containing EC and Hep, was reanalysed and a new unsupervised clustering was done to separate EC, Hep and doublets EC+Hep.

### Differential expression

The markers of each unsupervised cluster were identified with the R functions *FindAllMarkers with parameters only.pos = TRUE, min.pct = 0.25 and logfc.threshold = 0.25.* The markers of each time point were identified, with *Idents(x)* <-“*orig.ident*”, levels(Idents(x)) <-c(“0h”, “0h”, “3h”, “3h”, “6h”, “6h”, “24h”, “24h”), where *x* is the Seurat object followed by the R function *FindAllMarkers* with parameters *only.pos = TRUE, min.pct = 0.25* and *logfc.threshold = 0.25.*

### Stackplots

The percentage of cells in each cluster for each condition are displayed as stackplots with the R package *ggplot2*.

### Enrichment analysis

For individual gene lists, Pathway enrichment analysis was done with Metascape^32^ using as input statistically significant genes with log2FC > 0.25 with the option *Express Analysis*. For Multiple gene lists, pathway enrichment analysis was done with Metascape specifying as input the statistically significant genes with Log2FC > 0.25 with the option *Multiple Gene List* and *Express Analysis*.

### Cell Cycle

Cell cycle was assessed with the R function CellCycleScoring, with s.features and g2m.features provided by Seurat in the R object *cc.genes* after being transformed into mice genes with the R function *gorth* of the R package gprofiler2^33^.

### Dropout correction

For gene expression visualization, we used MAGIC^34^ to correct the dropout. First, a library size normalization was done with the R function *library.size.normalize* followed by the correction with the R function *magic* with parameters *genes=”all_genes”.*

### Violin plots

Violin plots were performed with the R function VlnPlot for the R package Seurat.

### Heatmap

MAGIC corrected data gene expression heatmaps were performed with heatmap.3 function from the R package heatmap3^35^. Each gene, x, was normalized with the function f(x)= x-min(x)/(max(x)-min(x)), and the data was clustered with average linkage clustering with Euclidean distance.

Average mRNA expression, from SCT transformed scRNA-seq expression, and average CyTOF expression heatmaps were performed with heatmap.2 from the R package gplots. Each gene, x, was normalized, f(x)= x-min(x)/(max(x)-min(x)) both for mRNA and Protein independently.

### Cell to cell communication

We assessed the cell communication with CellphoneDB 2^36^. Gene expression of every cell expression was normalized to UMI per 10000. Next, gene names were transformed into human gene names with the function *gorth* of the R package gprofiler2^33^, resulting duplicate gene names were removed and excluded from the analysis. Cells were grouped per time and cell type and inputed into cellphoneDB with *cellphonedb method statistical_analysis* with *options --threads=10*.

### Mass Cytometry

#### Cell staining

Cells were stained according to standard published methods for mass cytometry^37^. Briefly, 18 samples were viability stained for 30 minutes at 4°C with Cell-ID Cisplatin (Fluidigm, 201064) and barcoded with Cell-ID 20-plex Pd barcoding kit (Fluidigm, 201060) following the kit protocol. Barcoded samples were pooled and stained for 10 minutes at 4°C with anti-mouse CD16/32 Fc-block (eBiosciences, 14-0161-85). Cells were stained with the surface antibody cocktail (Extended Data Table 5) for 30 minutes at room temperature and then fixed and permeabilized with the eBioscience Foxp3/Transcription Factor Staining Buffer Set (ThermoFisher, 00-5523-00) following kit guidelines. The intracellular/intranuclear staining cocktail (Extended Data Table 5) was then added in the kit permeabilization buffer for 90 minutes at 4°C. Cells were washed in MACS buffer (PBS + .7% FBS + 2mM EDTA) and placed in Maxpar Fix & Perm Buffer (Fluidigm, 201067) containing Cell-ID Intercalator-Ir (Fluidigm, 201192B). The sample was kept at 4°C until acquisition the following day on a Helios mass cytometer at the Cytometry Facility at the University of Zürich.

### CyTOF data preprocessing

FCS files were pre-processed using the premessa R package for pre-processing of flow and mass cytometry data by Pier Federico Gherardini at Parker Institute for Cancer Immunotherapy. In premessa, we concatenated our FCS files and de-barcoded using the values of .2 Minimum separation and 30 Maximum Mahlanobis distance. The 12 FCS files were then uploaded into Cytobank and target expression was checked overtime by plotting dot plots with all Panel/Channel values vs Time. No pressure drops or spikes were observed over time and the positive signals were well separated and stable. Live/single cells were gated on by plotting DNA (191Ir) vs Cisplatin (198Pt) and gating on the DNA positive, Cisplatin negative cells and tailoring the gates as needed for each of the samples. After, Pairwise Plots were checked in Cytobank for compensation issues/debris and a debris signal was gated out by gating on 139La vs 140Ce negative cells. The resulting FCS files were imported into R and analyzed with the CATALYST R package for Cytometry dATa anALYSis Tools^38^. Briefly, 12 FCS files were read as a flowSet object with the R function *read.flowSet*. The flowSet was added into a Single Cell Experiment (SCE) object containing the panel data (targets and channels used) and the sample metadata (conditions, sample ID, patient ID) CSV files. The formation of the SCE object with *prepData* also arcsinh transformed the data with a cofactor of 5. We checked the quality of the samples by checking the number of cells per sample with *plotCounts*, and by looking for outliers with the multiple-dimensional scaling (MDS) *plotMDS*, and by plotting the marker expression per sample with *plotExprHeatmap*.

### CyTOF Clustering

The data was clustered with the R function *cluster* in CATALYST, which is a wrapper that performs high resolution FlowSOM clustering and lower resolution ConsensusClusterPlus metaclustering. By default the data is initially clustered into 100 groups (xdim = 10, ydim = 10) and then the function metaclusters to a default of 20 clusters (maxK = 20). We selected the lineage markers (CD3, CD45, Ly6G, CD11c, F4/80, CD19, Ly6C, GATA6, CD206, and CD11b) and clustered the data into 25 clusters with a seed of 1. We plotted these clusters with *plotClusterHeatmap* to see the target expression in each cluster and identify the different cell populations present. Once we identified the populations in the liver, we made a merging table CSV and used *mergeclusters* to manually merge the 25 clusters into the 9 main cell populations in the liver (T cells, B cells, Hepatocytes, KC, BMDM, Granulocytes, LSECs, pDCs, and cDCs). Later in the analysis, we used *filterSCE* to select the BMDM cells and performed clustering on these with maxK = 7 and seed = 1.

### CyTOF Dimensionality reduction

t-SNE dimensionality reduction was done using all panel targets with the R function *runDR* in CATALYST with the parameters *dr = “TSNE”* and for *cells* = 1000 for 1,000 cells per sample. We then filtered the SCE object with *filterSCE* to select only the macrophage population (BMDM) and performed dimensionality reduction with t-SNE using a *perplexity = 10*, and the minimal cell number available for this population across all samples *(cells = min(n_cells(x))*.

### CyTOF Differential Abundance

The differential abundance (DA) of each liver subset over time was assessed with the *diffcyt* package^39^. *Diffcyt* provided us with statistical methods for differential discovery over time. This is done by first specifying a design matrix based on the samples and conditions that will be tested with *createDesignMatrix*. A contrast matrix is then created to specify the contrast being testing (the control samples compared to each timepoint) with *createContrast*. For the comparison of 0 hours with all time points after PHx, the contrast was the vector (−1,1/3,1/3,1/3). For comparison of 0 hours and 3 hours with 6 and 24 hours, the contrast was the vector (−0.5, −0.5, 0.5, 0.5). *diffcyte* uses the *diffcyte-DA-edgeR* method for the DA analysis and the DA heatmap was displayed with the R function *PlotDiffHeatmap*.

### Barplots differential abundance

Each sample was normalized by its number of cells. Second, the fraction of each cell type was divided by its maximum over time. Third, the two replicates of scRNA-seq were averaged, and the three replicates of CyTOF were averaged at each time point.

### Flow Cytometry

To label the nascent DNA in vivo, 5-ethynyl-2’-deoxyuridine (EdU) was injected intraperitoneally 24 hours post PHx. The dosage of EdU used was 100mg/kg. After two hours, the liver was perfused and digested in situ as described above. After liver digestion, cells were resuspended in MACS buffer (PBS + .7% FBS + 2mM EDTA). Cells were centrifuged at 50g for 5 minutes at 4°C and the supernatant was collected and the hepatocyte pellet was discarded. After resuspension in PBS, cells were counted and 2.5 million cells were used for staining.

### Cell subset staining

(all incubations done at 4°C in the dark)

Viability staining was performed in PBS with eBioscience Fixable Viability Dye eFluor 506 (ThermoFisher, 65-0866-14) at 1:1000 for 30 minutes. FC receptors were blocked with FC-block at 1:1000 for 10 minutes. Surface staining was done by adding the flow cytometry antibody cocktail outlined in Extended Data Table 6 containing F4/80 (BioLegend, 123120), CD11b (BioLegend, 101222), CD19 (BioLegend, 302218), CD45 (BD Biosciences, 564279), CD31 (BioLegend, 102528), CD3 (BioLegend, 100330), NK1.1 (BioLegend, 108724), CD206 (BioLegend, 141706) for 15 minutes.

### Detection of EdU incorporation

ClickiT EdU Alexa Fluor 647 Flow Cytometry Assay (ThermoFisher, C10419) kit was used to detect EdU incorporation into nascent DNA, and DNA content was assessed by adding a drop of FxCycle Violet Stain (ThermoFisher, F10347) to each sample a half hour before acquisition on the Fortessa flow cytometer.

### Flow cytometry data analysis

FCS files were loaded into FlowJo10 version 6.11 and cells were cleaned with a standard gating strategy for non-hepatocytes (FSC-A vs SSC-A), single cells (FSC-A vs FSC-H), living cells (BV506 vs F4/80) and lymphocytes (CD45+) as show in Supplemental Figure 23 Cells that were not of interest (T cells, B cells, NK cells, endothelial cells) were removed by gating on dump channel negative cells (CD3, CD19, CD31, NK1.1), followed by Kupffer cell identification with high expression of F4/80 and low expression of CD11b. Within Kupffer cells, the percentage of EdU positive cells is shown with quadrant gates on a log scale and DNA content on a linear scale in Figure 3c.

### Spinning disk confocal intravital microscopy (SD-IVM)

Intravital imaging experimental animal protocols were approved by the University of Calgary Animal Care Committee and were compliant with the Canadian Council for Animal Care Guidelines. WT C57BL/6J were purchased from The Jackson Laboratory. Clec4f-cre-tdTomato mouse (kind gift from Cristopher K. Glass, UCLA)^40^ were bred to the B6.Cg-Gt(ROSA)26Sortm6(CAG-ZsGreen1)Hze/J (Jackson) the offspring of these mice express tdTomato located in the nucleus and ZS-green in the cytosol under direction of the C-type lectin domain family 4, member f (Clec4f) promoter in Kupffer cells. All animals were maintained in a specific pathogen–free environment at the University of Calgary Animal Resource Centre. Mice were housed under standardized conditions of temperature (21– 22°C) and illumination (12 h light/12 h darkness) with access to tap water and pelleted food ad libitum. Antibodies against F4/80 (BM8) was obtained from eBioscience. A tail vein catheter was inserted into mice after anesthetization with 200mg/kg ketamine (Bayer Animal Health) and 10mg/kg xylazine (Bimeda-MTC). Partial hepatectomy was performed described above. Surgical preparation of the liver remnant was performed as previously described^41^, however, due to the resection of left liver lobe, the right lobe was prepared for imaging. Mouse body temperature was maintained at 37°C with a heated stage. Image acquisition was performed using Olympus IX81 inverted microscope, equipped with an Olympus focus drive and a motorized stage (Applied Scientific Instrumentation) and fitted with a motorized objective turret equipped with 10x/0.40 UPLANSAPO and 20x/0.70 UPLANSAPO objective lenses and coupled to a confocal light path (WaveFx; Quorum Technologies) based on a modified Yokogawa CSU-10 head (Yokogawa Electric Corporation). Target cells were visualized using fluorescent reporter cells. In addition, KCs were stained by i.v. administration of 2.5µg anti-F4/80 fluorescent conjugated monoclonal Antibody. Laser excitation wavelengths 491, 561, 642, and 730 nm (Cobolt) were used in rapid succession, together with the appropriate band-pass filters (Semrock). A back-thinned EMCCD 512 × 512 pixel camera was used for fluorescence detection (Hamamatsu). Volocity software (Perkin Elmer) was used to drive the confocal microscope. Imaris software was used for 3D rendering and analysis of images.

### Phxatlas.ch data visualization

Dropout corrected gene expression was uploaded to phxatlas.ch webpage. Each gene can be visualized as a tSNE plot of the dropout corrected expression. Boxplot of the dropout corrected expression in each sample and separated cell type.

## Extended Data Figure Legends

**Extended Data Fig. 1.**
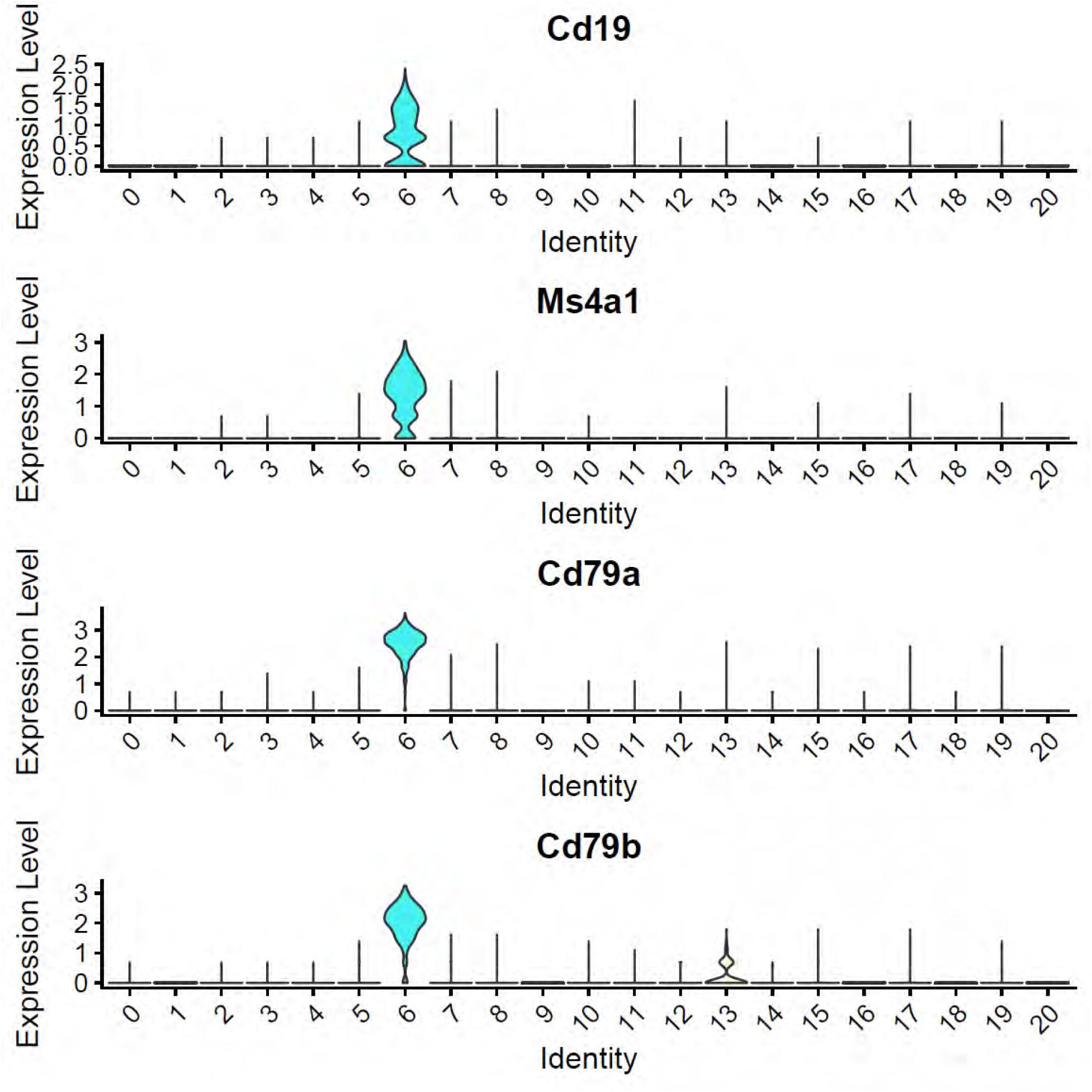
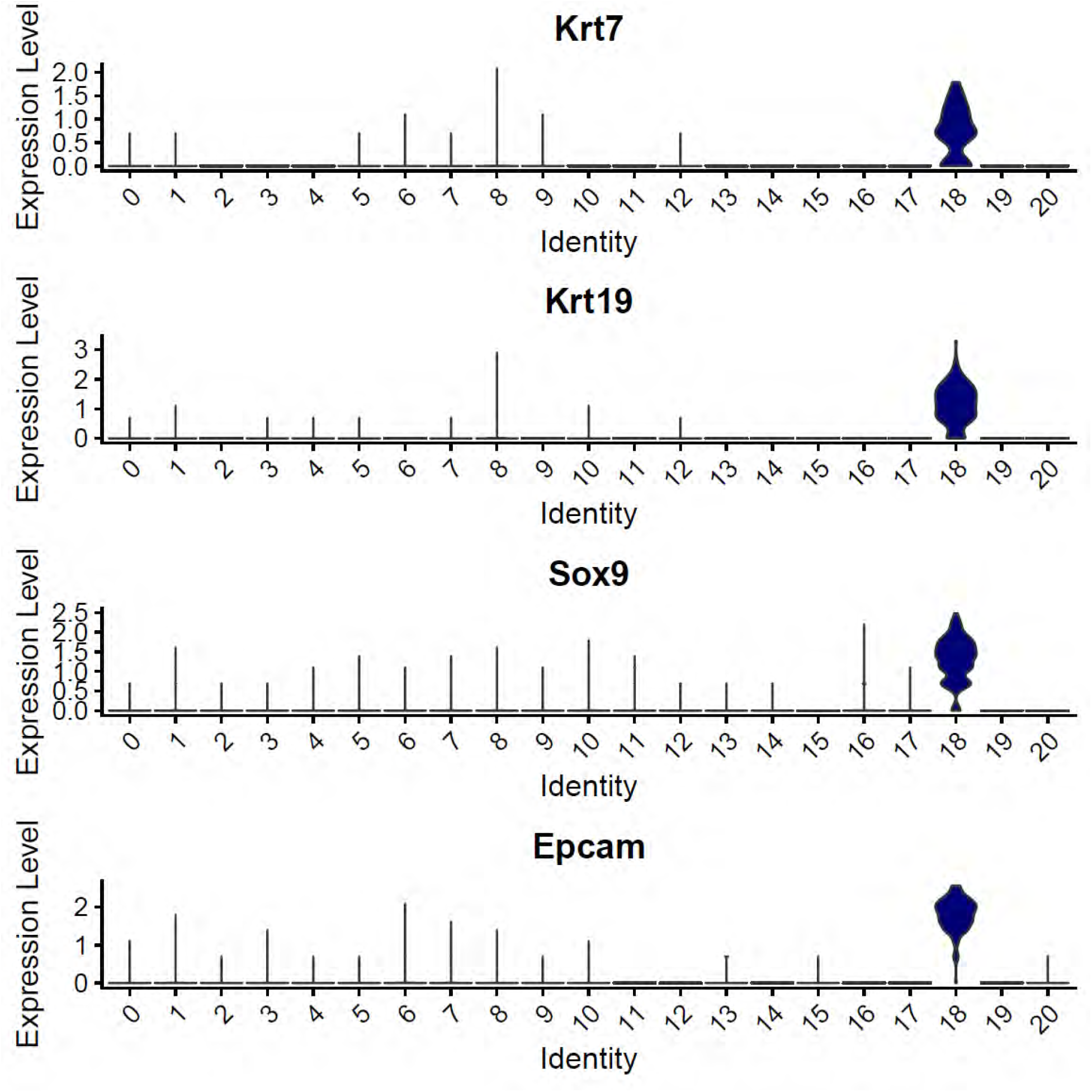
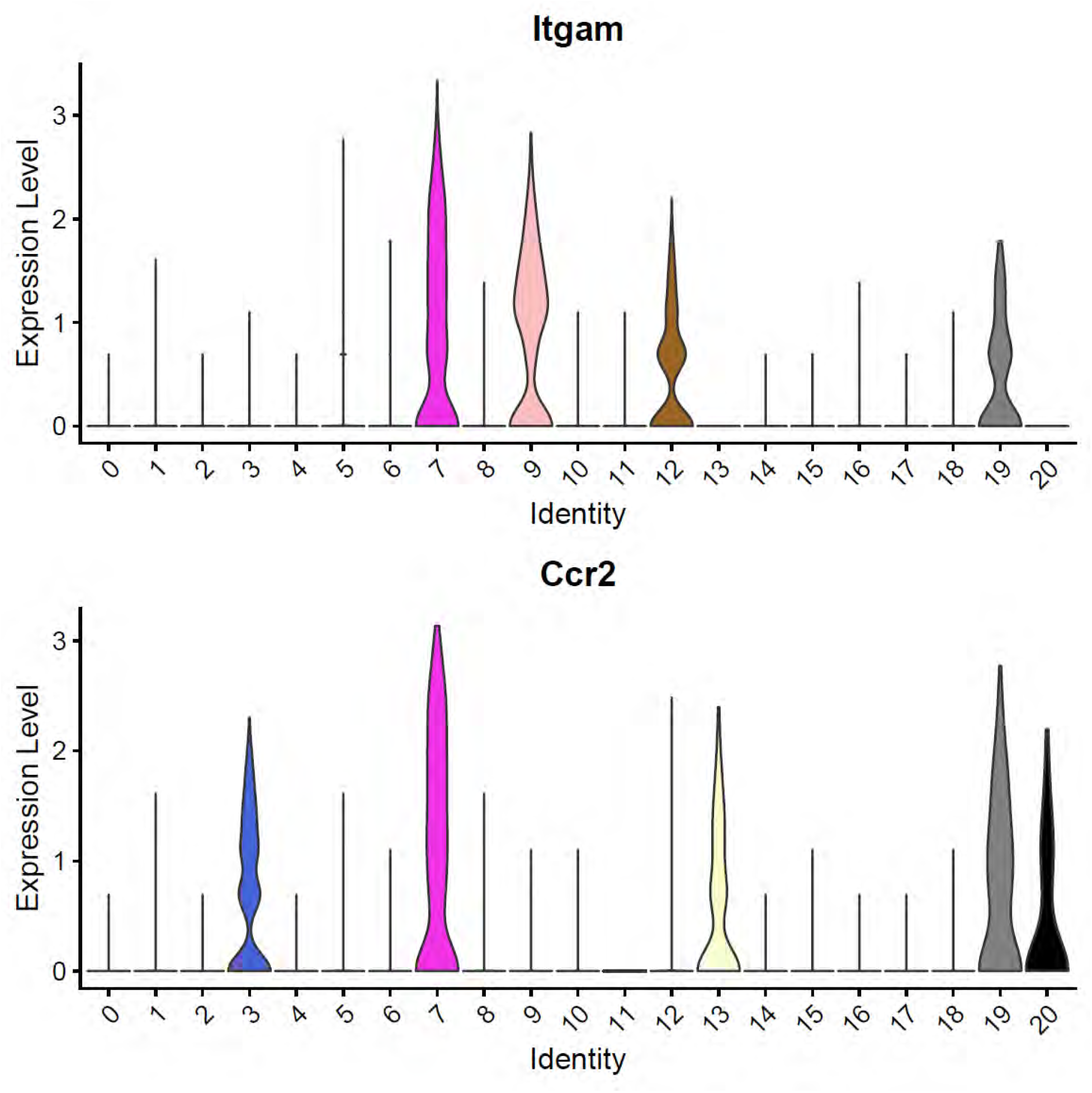
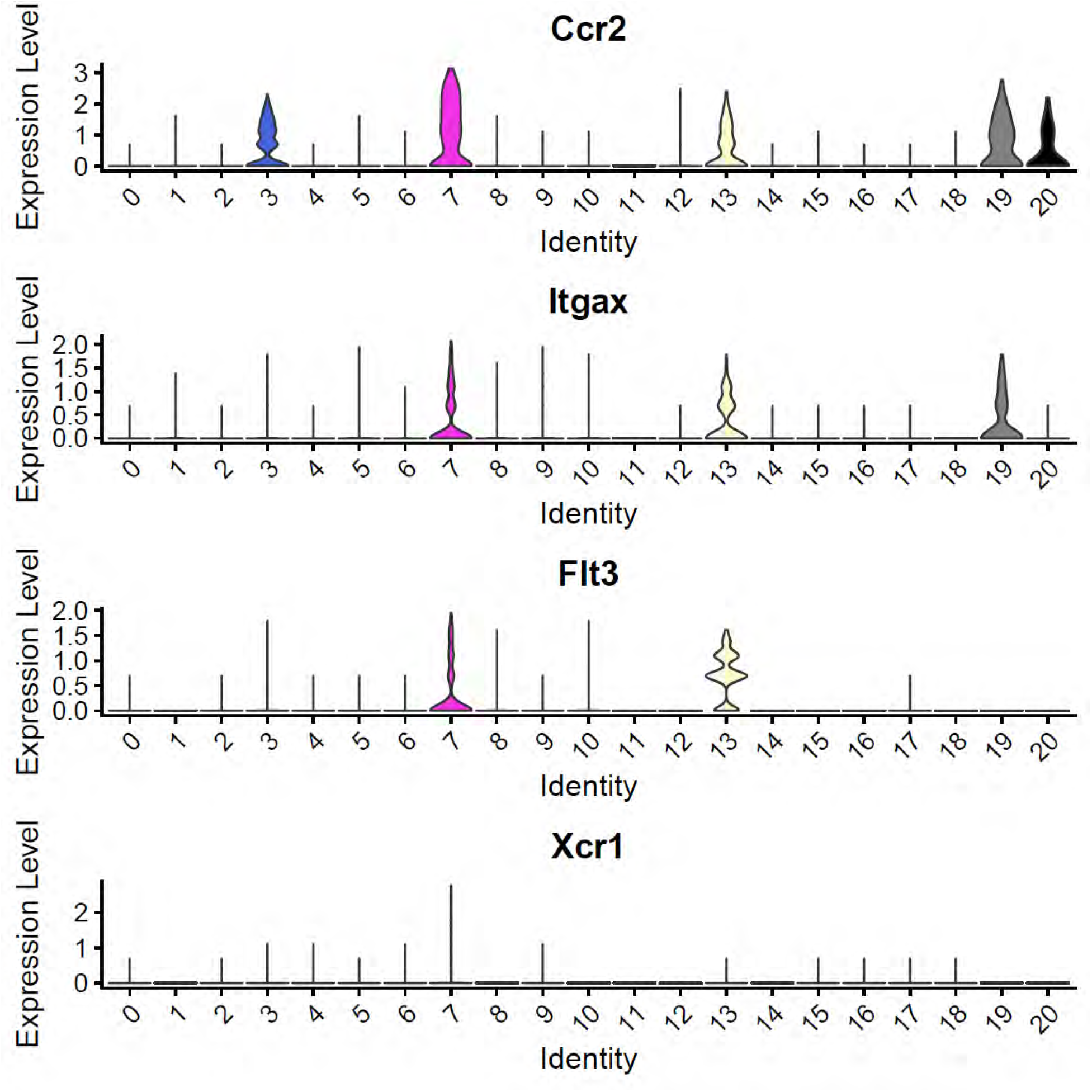
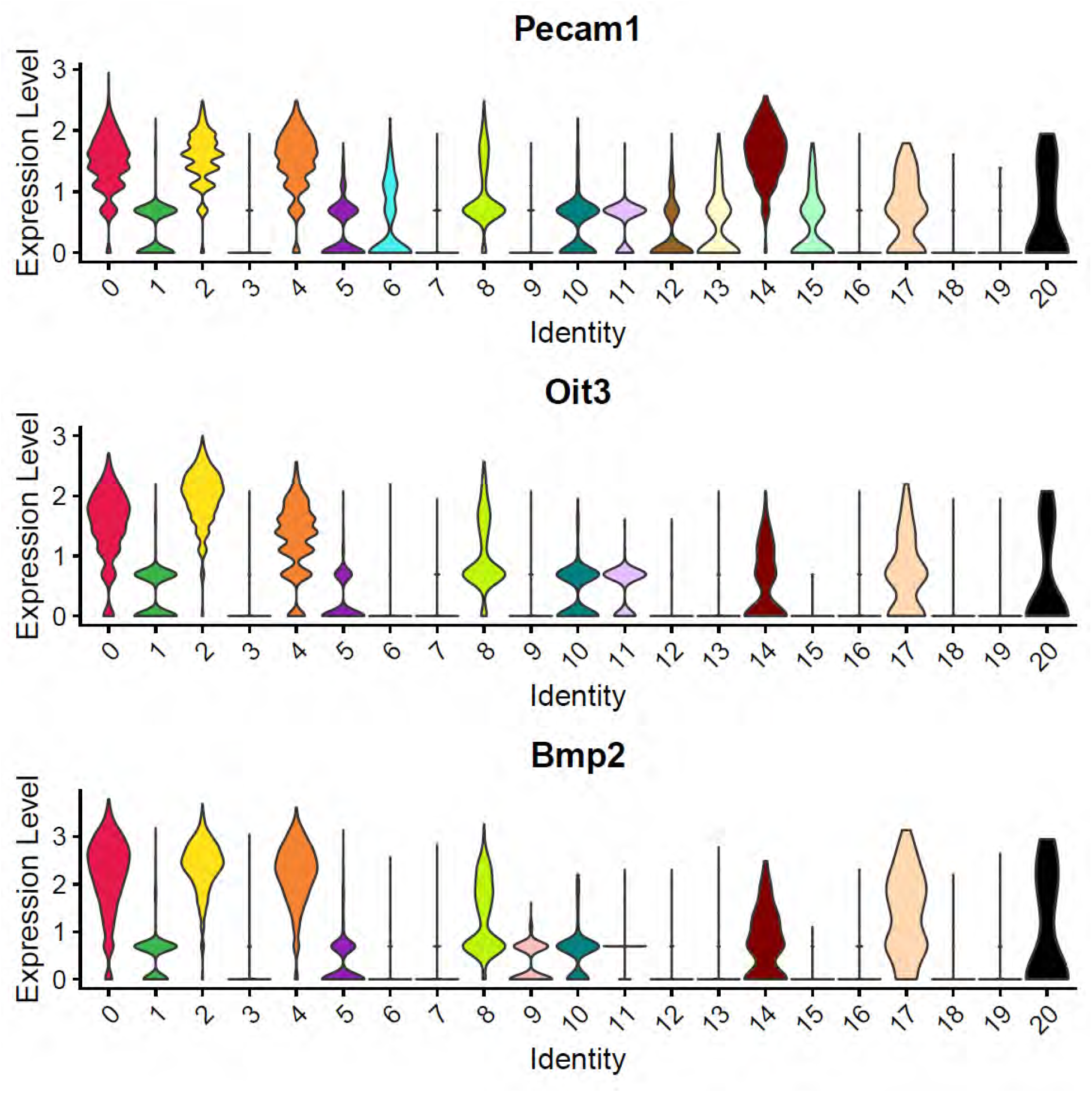
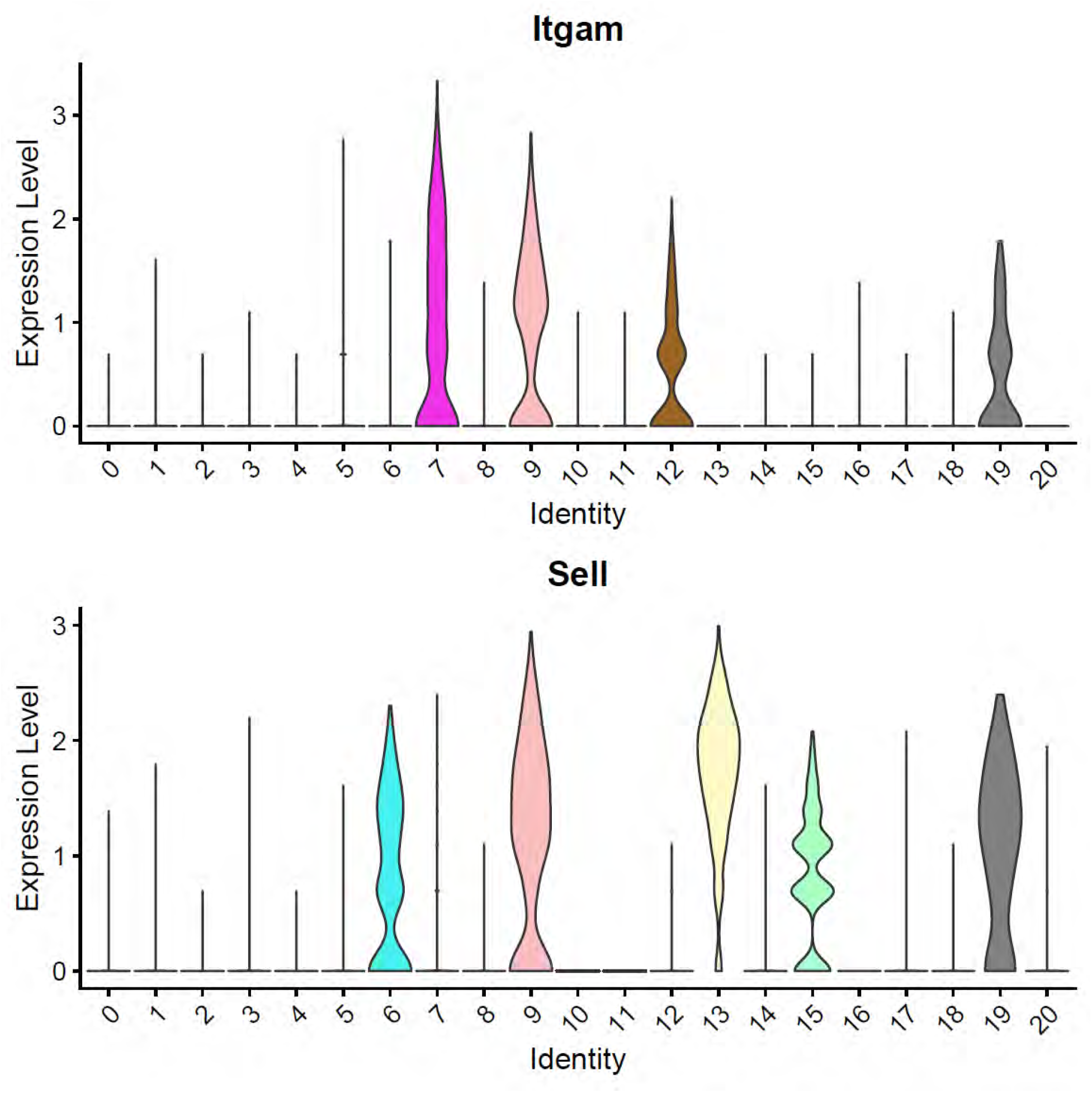
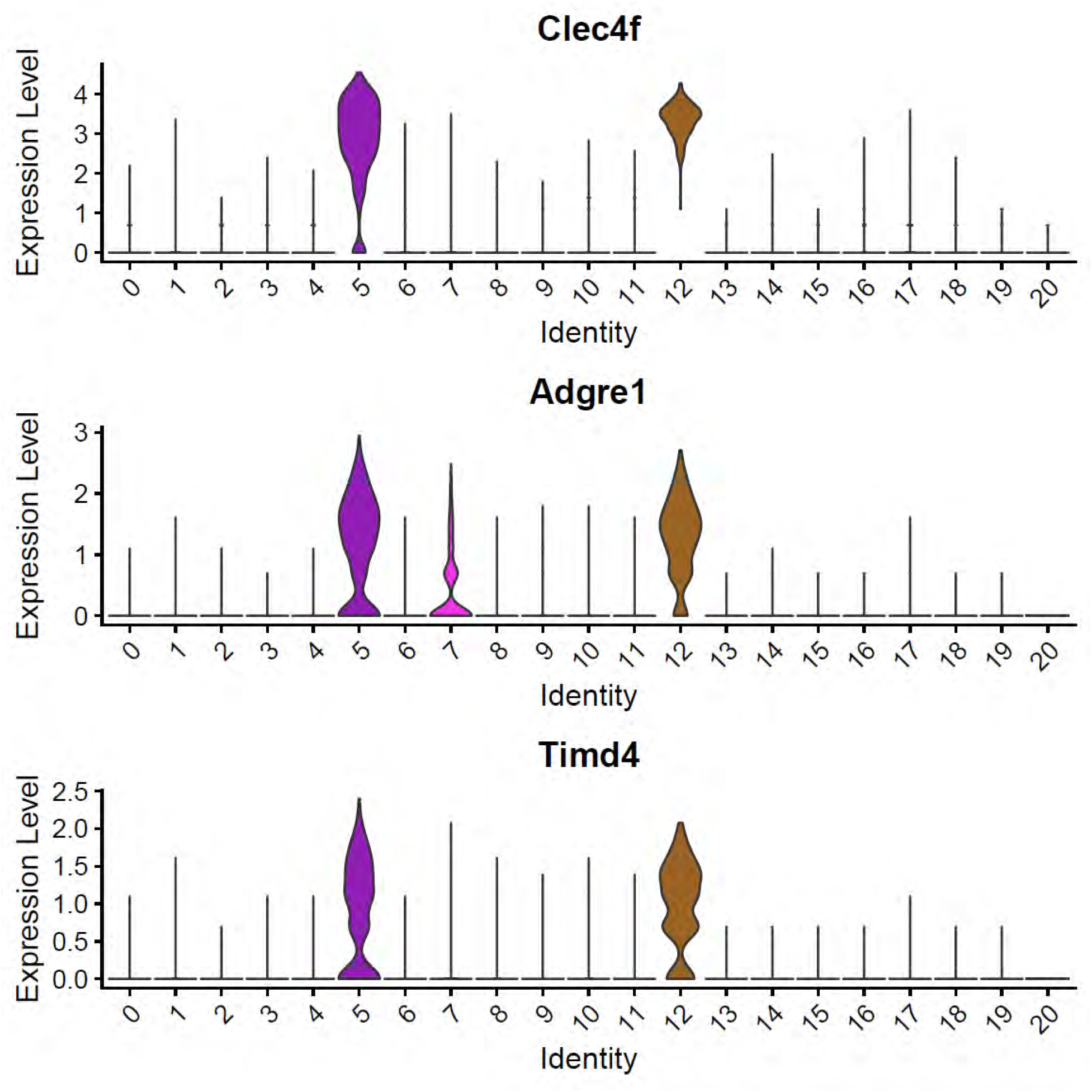
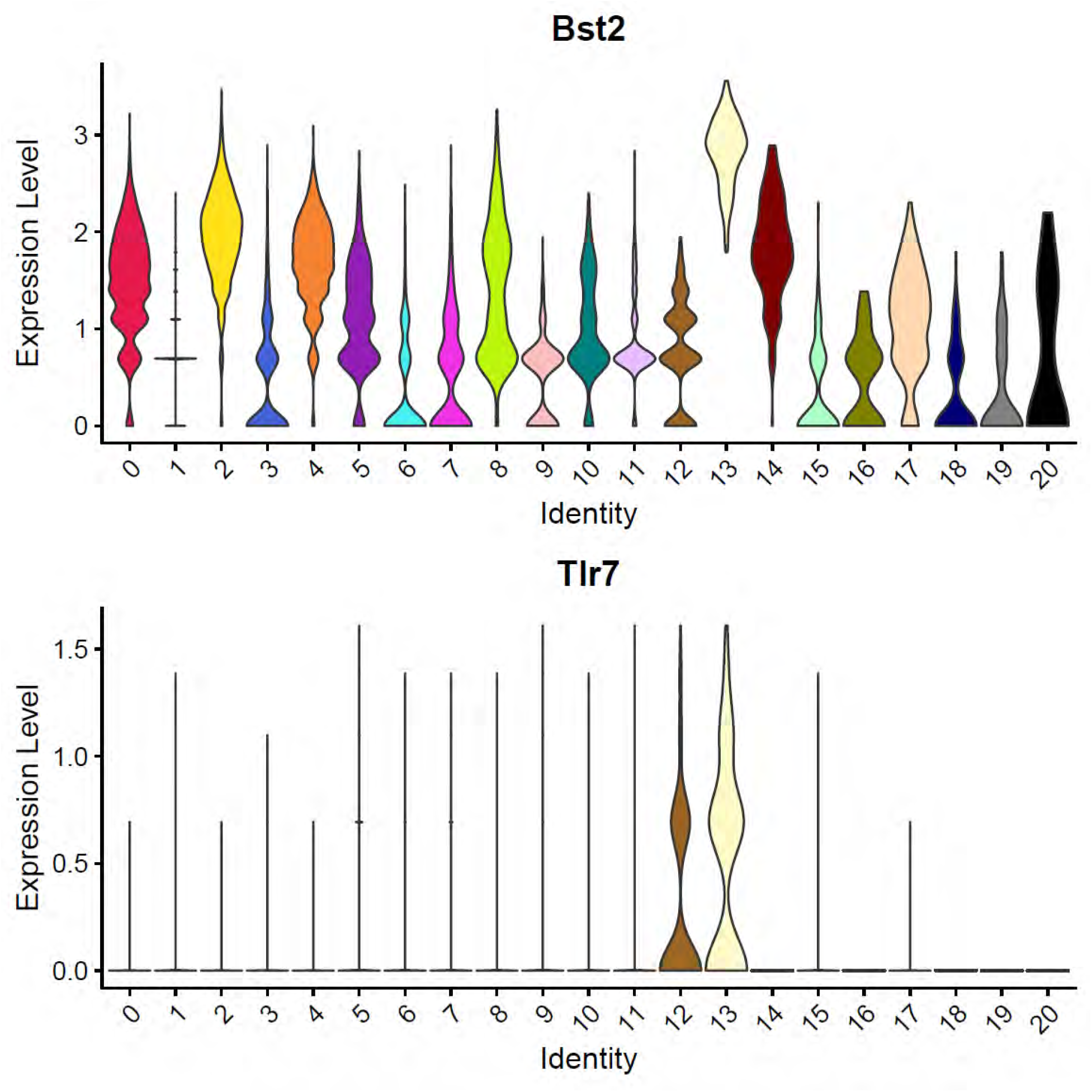
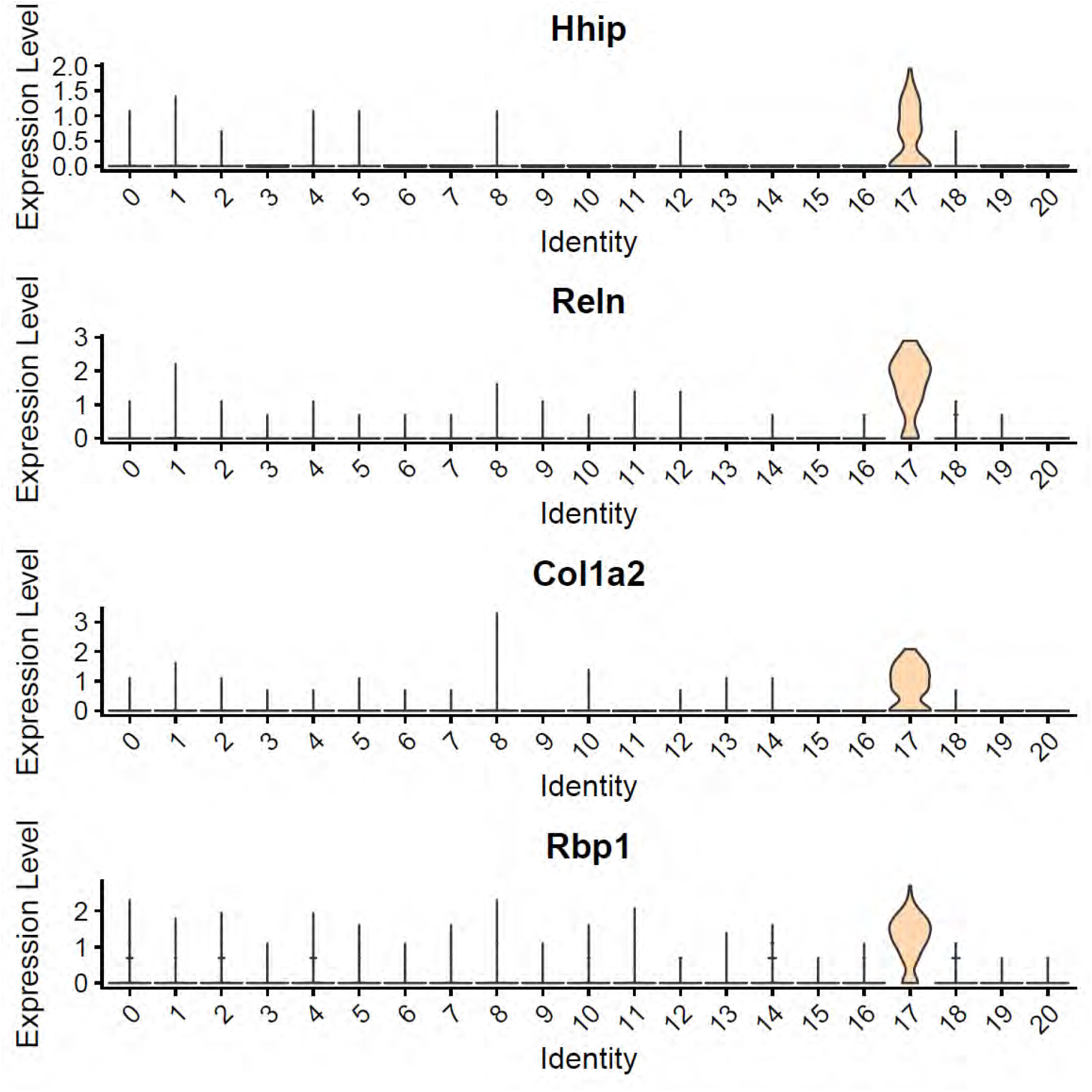
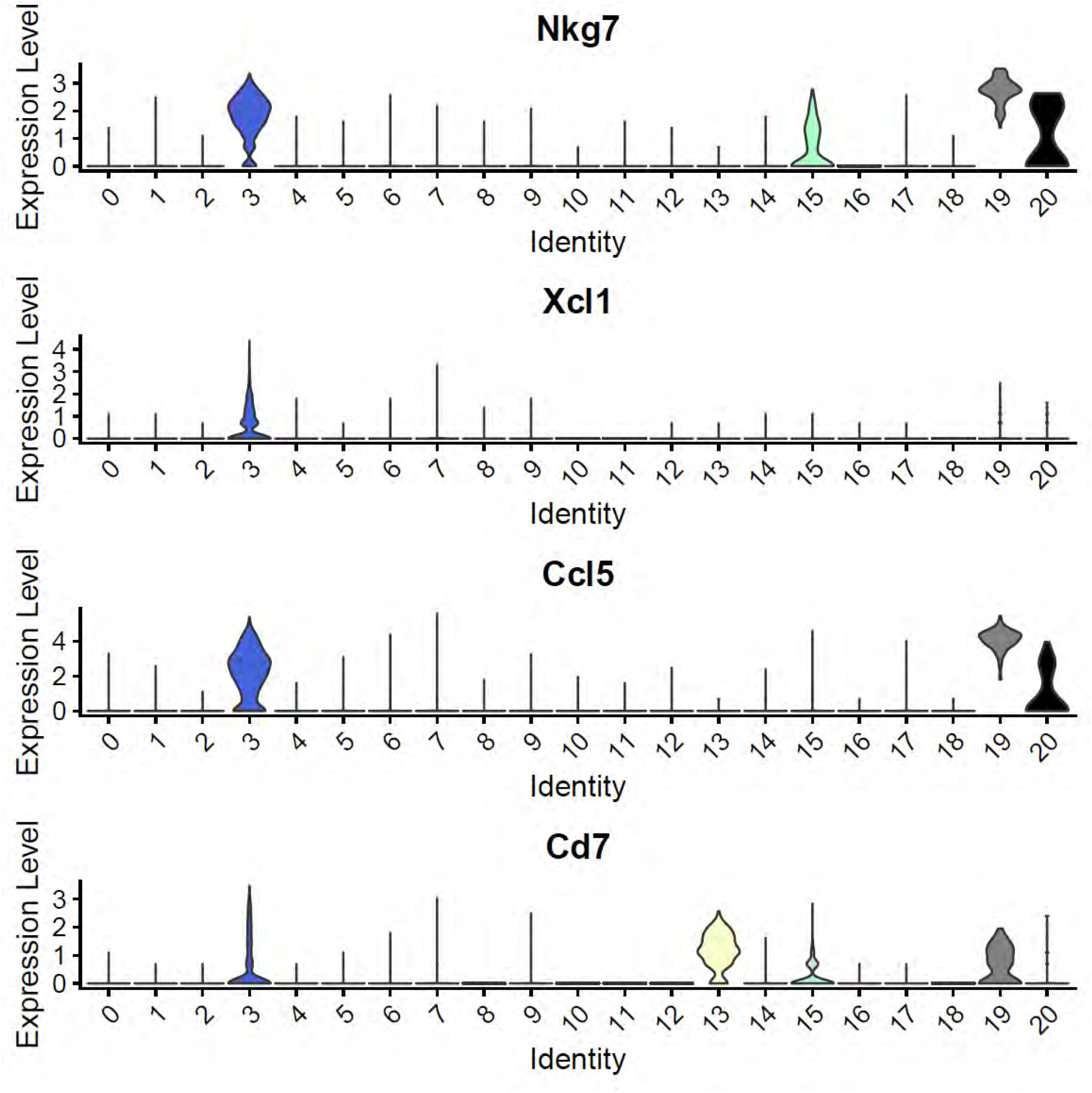
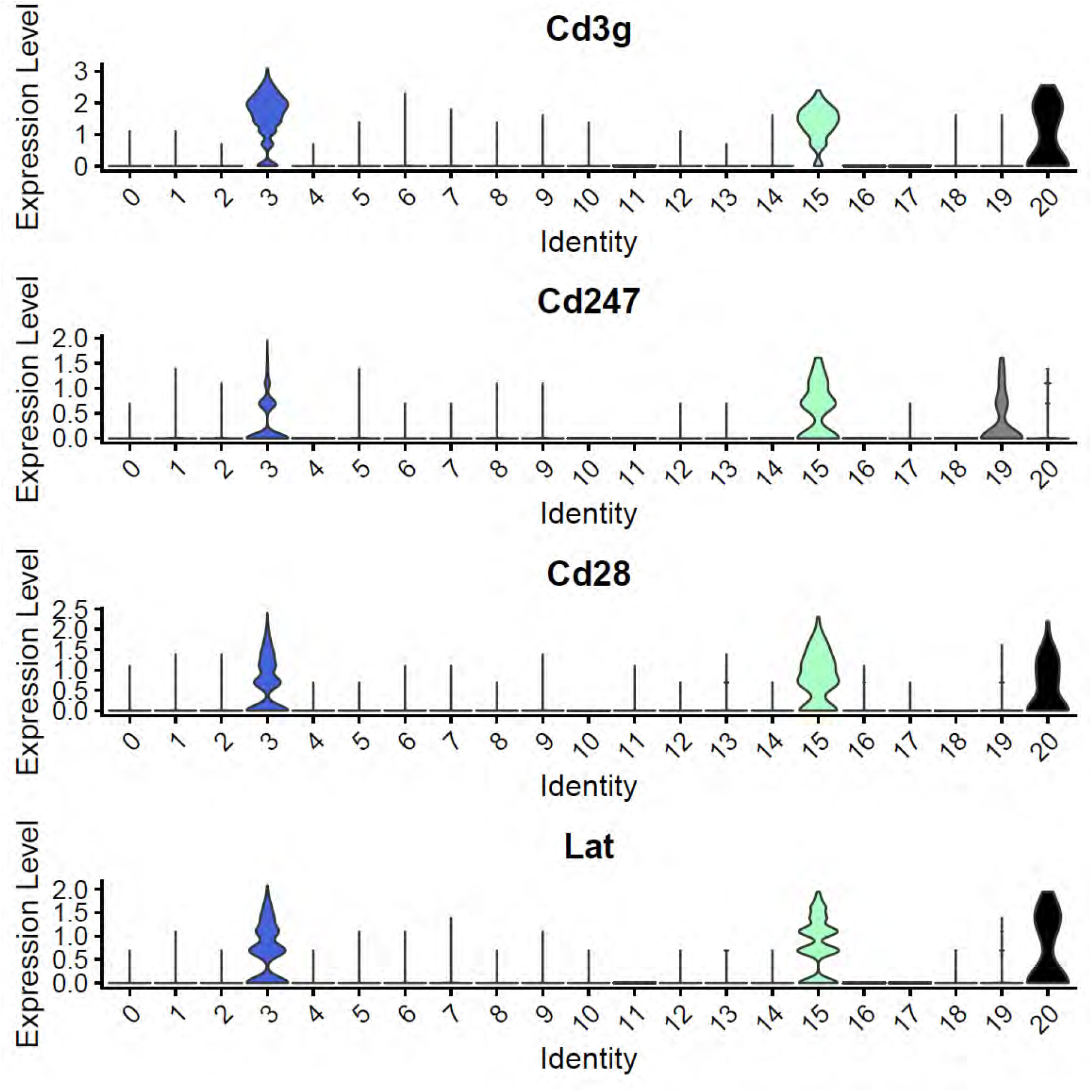
Marker expression. Violin plots showing the gene expression of markers used to identify each cell population present in the liver (see table T2). Cell subsets were identified as follows: B cells (6), biliary epithelial cells (18), BMDM and classic dendritic cells (7) later split (see Extended Data Fig 2), endothelial cells (0, 2, 4, 14), granulocytes (9), hepatocytes (1, 10, 11, 16), Kupffer cells (12), plasmacytoid dendritic cells (13), stellate cells (17), T cells (3, 15, 19), unknown (20). Additionally Cluster 5 seem to contain hepatocytes, Kupffer cells or doublets and Cluster 8 seem to contain endothelial cells, hepatocytes or doublets, and were studied later on (see Extended Data Fig 3).

**Extended Data Fig. 2.**
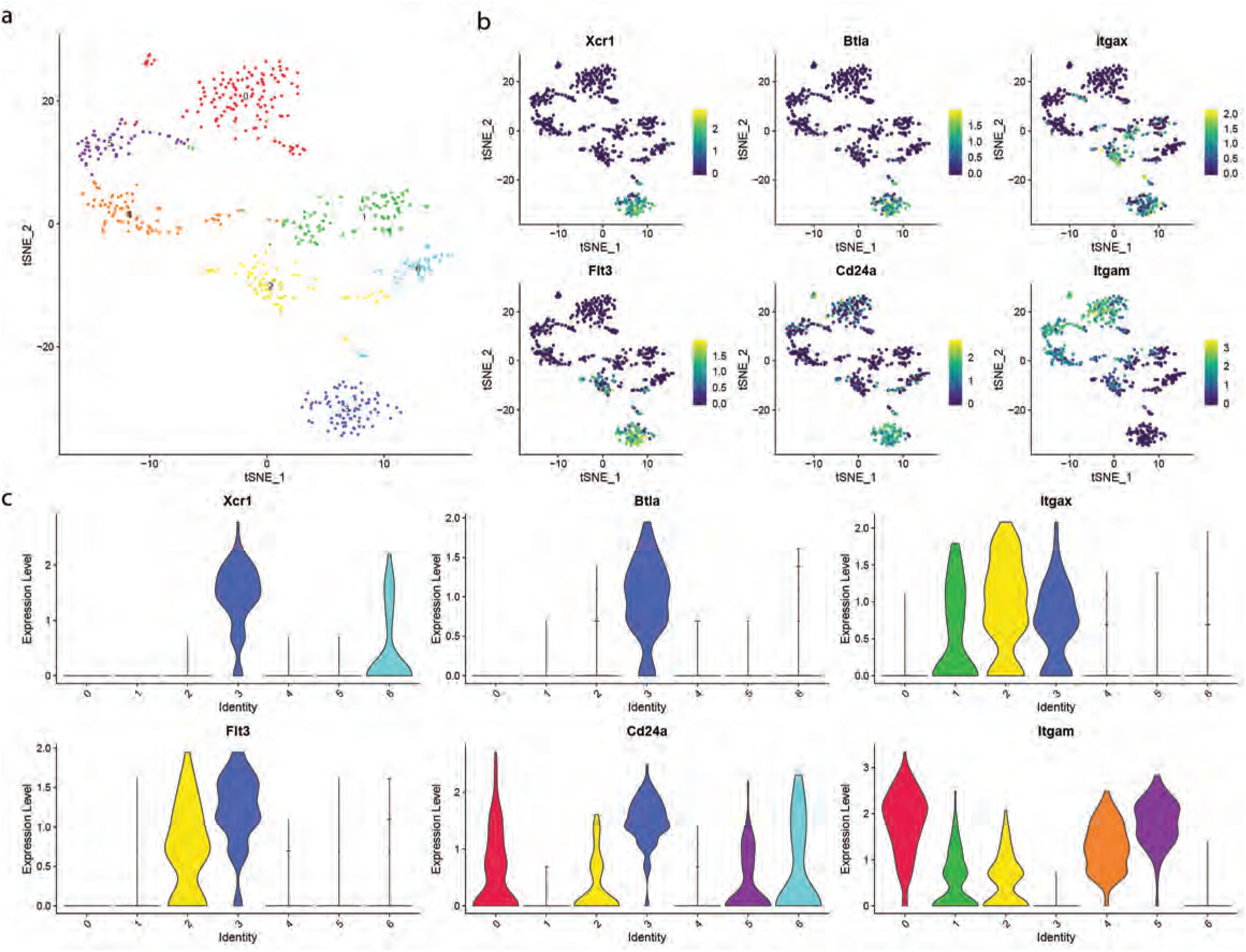
cDC identification. cluster 7 a) t-SNE colored by unsupervised cluster, b) t-SNE colored by classic dendritic cells marker expression, c) violin plot of the marker expression colored by cluster. Cells were identified as follows: BMDM (0,1,2,4,5), classic dendritic cells (3).

**Extended Data Fig. 3.**
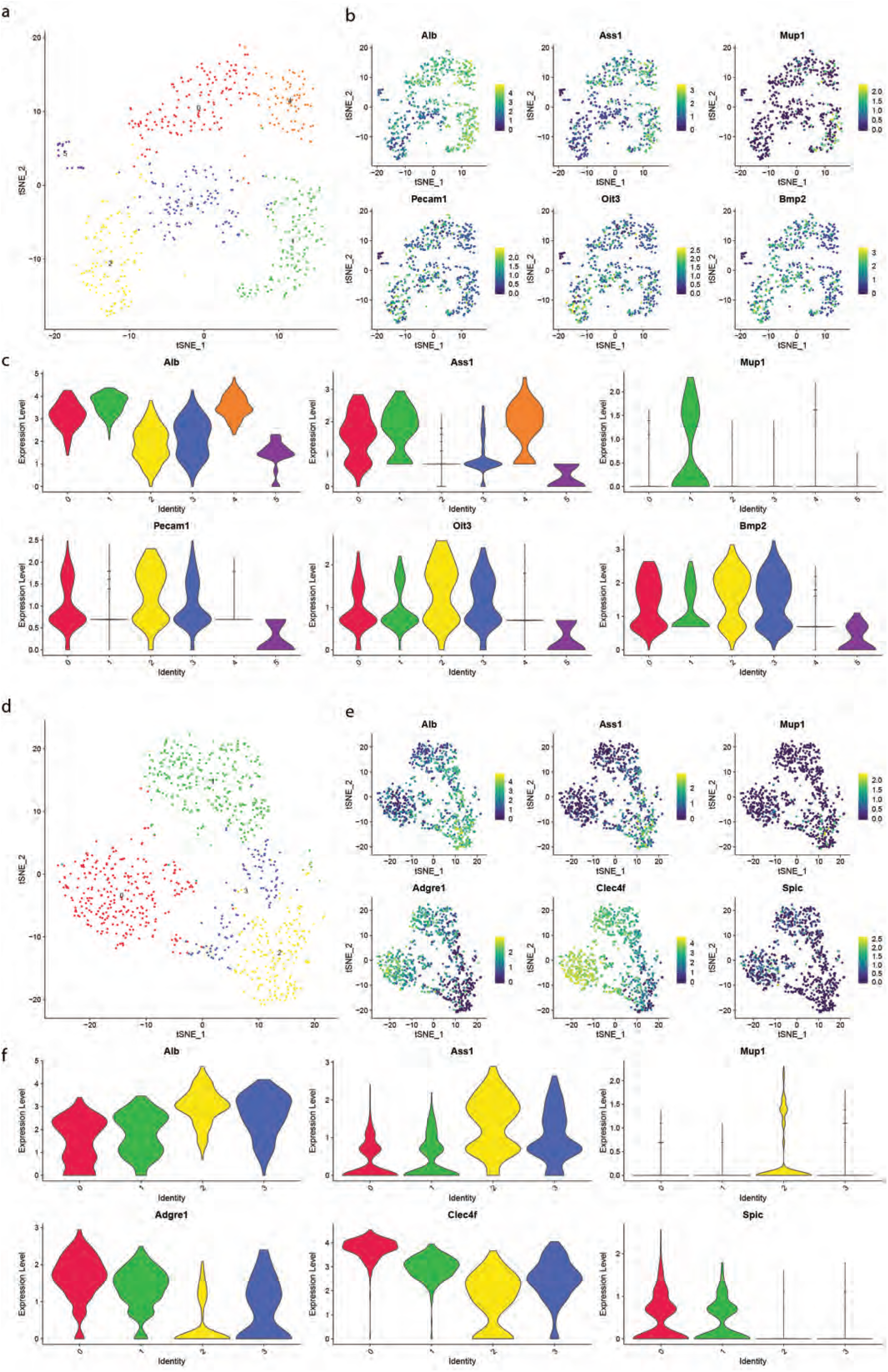
Identification of doublets. cluster 8 a) t-SNE colored by unsupervised cluster, b) t-SNE colored by endothelial cells and hepatocyte marker expression, c) violin plot of the marker expression colored by cluster. Cells were identified as follows: hepatocyte (1,4), endothelial cells C (2,3), doublets hepatocytes+Kupffer cells (0,5). Cluster 5 d) t-SNE colored by unsupervised cluster, e) t-SNE colored by hepatocyte and Kupffer cell marker expression, f) violin plot of the marker expression colored by cluster. Cells were identified as follows: Kupffer cells (0,1), doublets hepatocytes+Kupffer cells (2,3).

**Extended Data Fig. 4.**
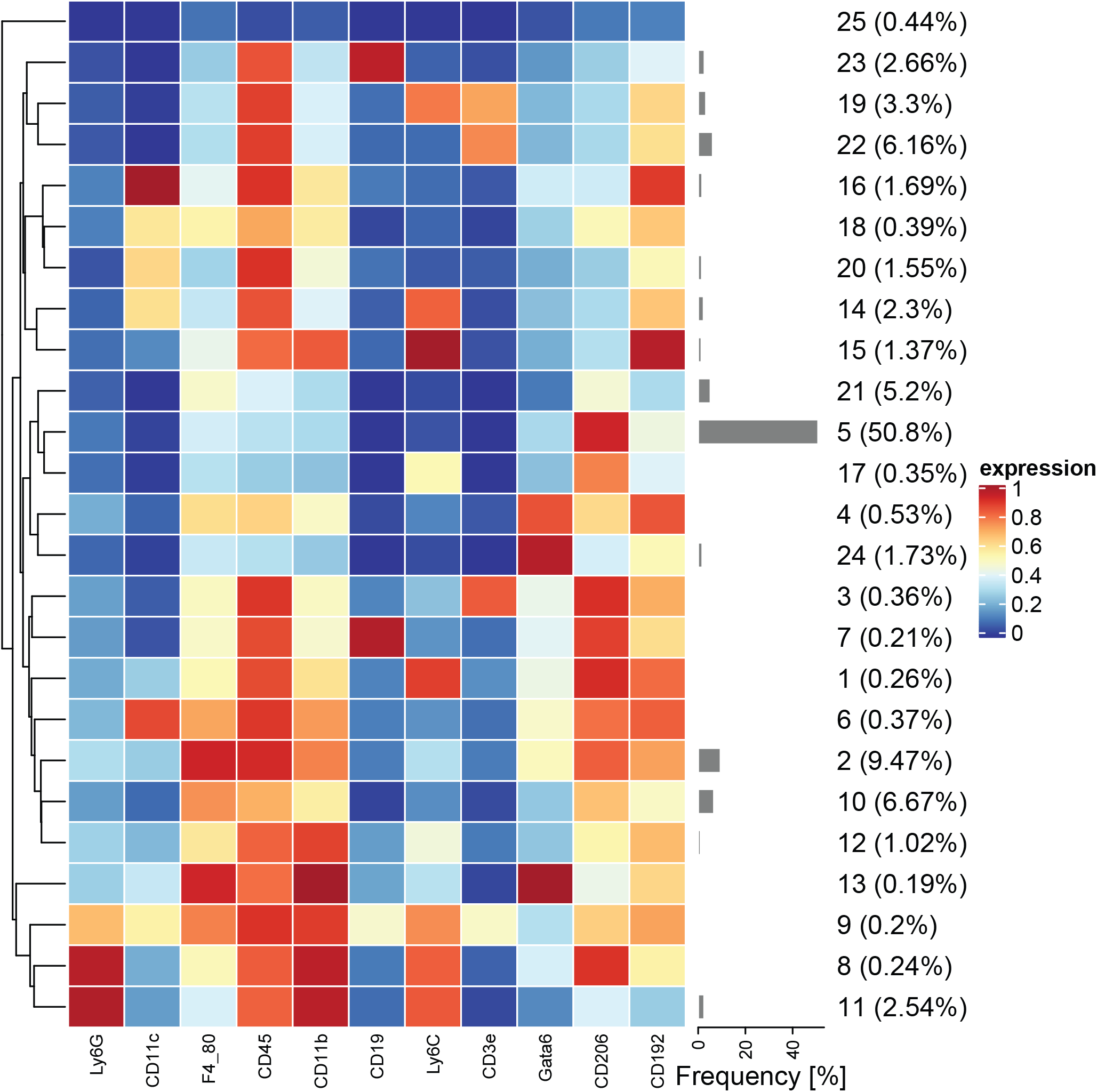
CyTOF Classification. Heatmap of target expression and cluster frequencies of the 25 flowsom unsupervised clusters. Cell subsets were identified as follows: B cells (23, 7), BMDM (6, 1, 4, 13, 15, 12, 20, 18), classic dendritic cells (16), endothelial cells (5, 17), granulocytes (8, 11), hepatocytes (24), Kupffer cells (2, 10, 21), plasmacytoid dendritic cells (14), T cells (19, 22, 3), unknown (25, 9).

**Extended Data Fig. 5.**
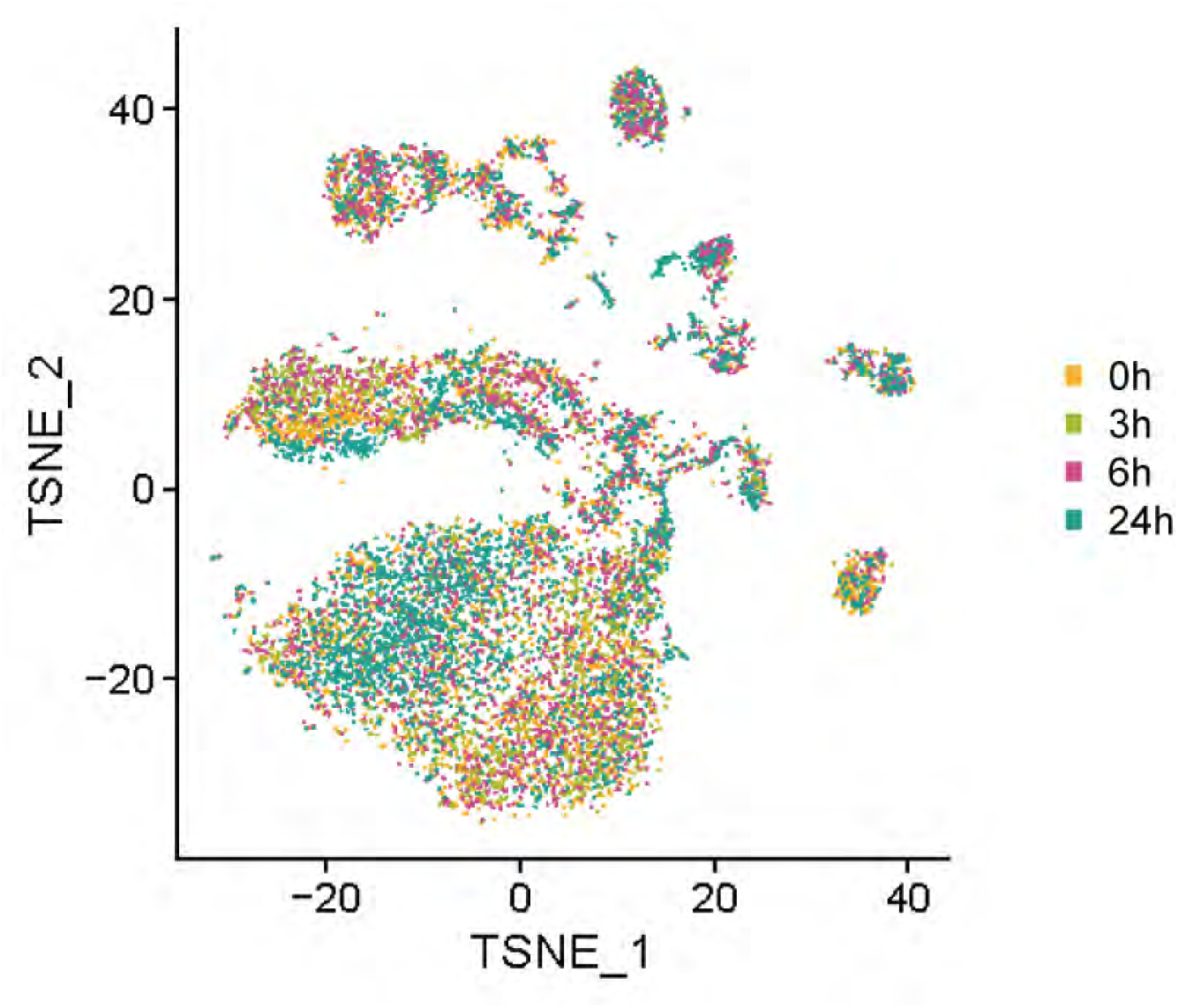
t-SNE visualization of CyTOF colored by sample.

**Extended Data Fig. 6.**
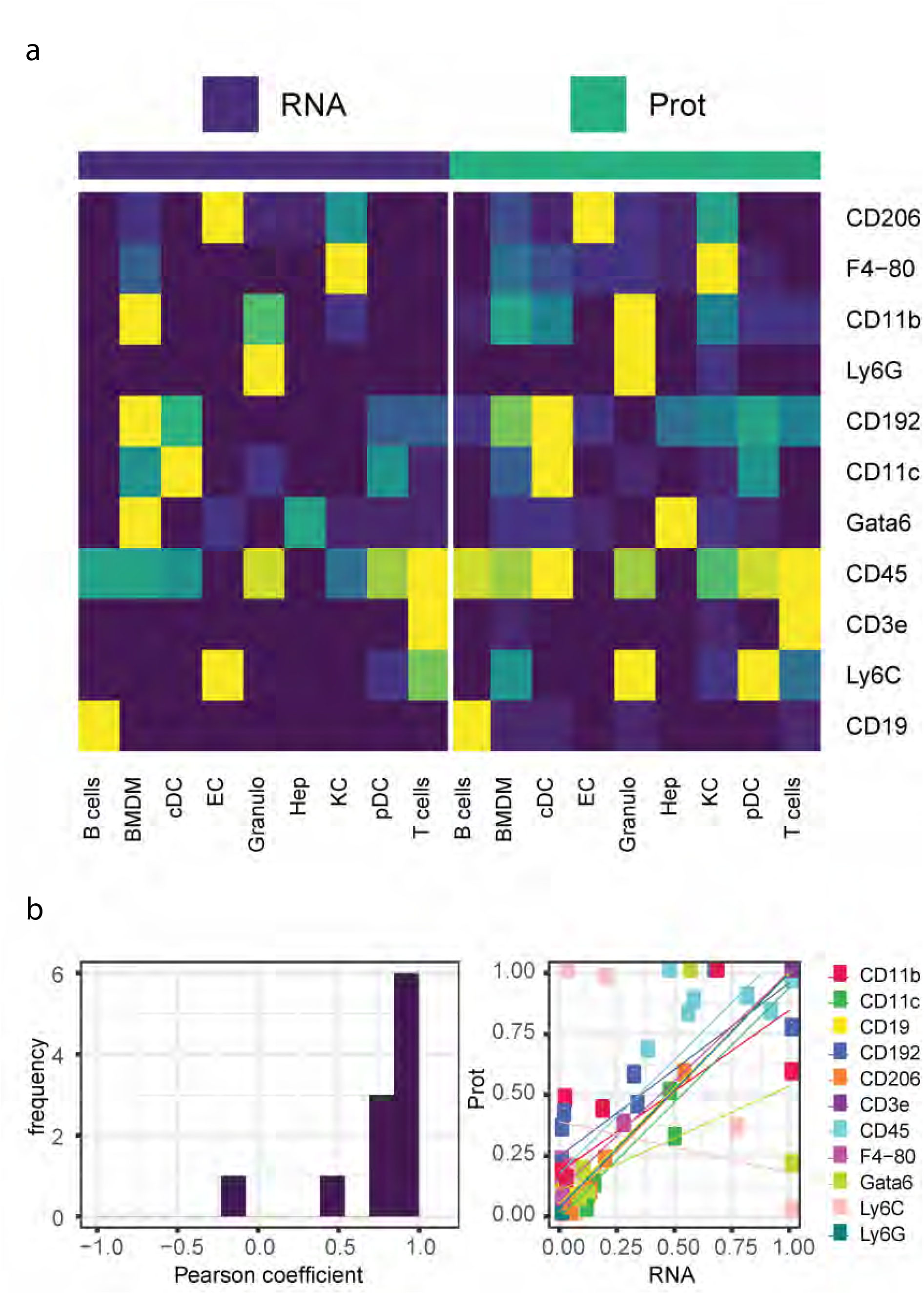
a) Heat map of average mRNA (purple) and protein (green) expression in each cell type used for CyTOF lineage markers. b) (left) Distribution of Pearson correlation of mRNA and protein expression of the lineage markers, (right) scatter plot and linear regression of the normalized mRNA and protein expression of the lineage markers.

**Extended Data Fig. 7.**
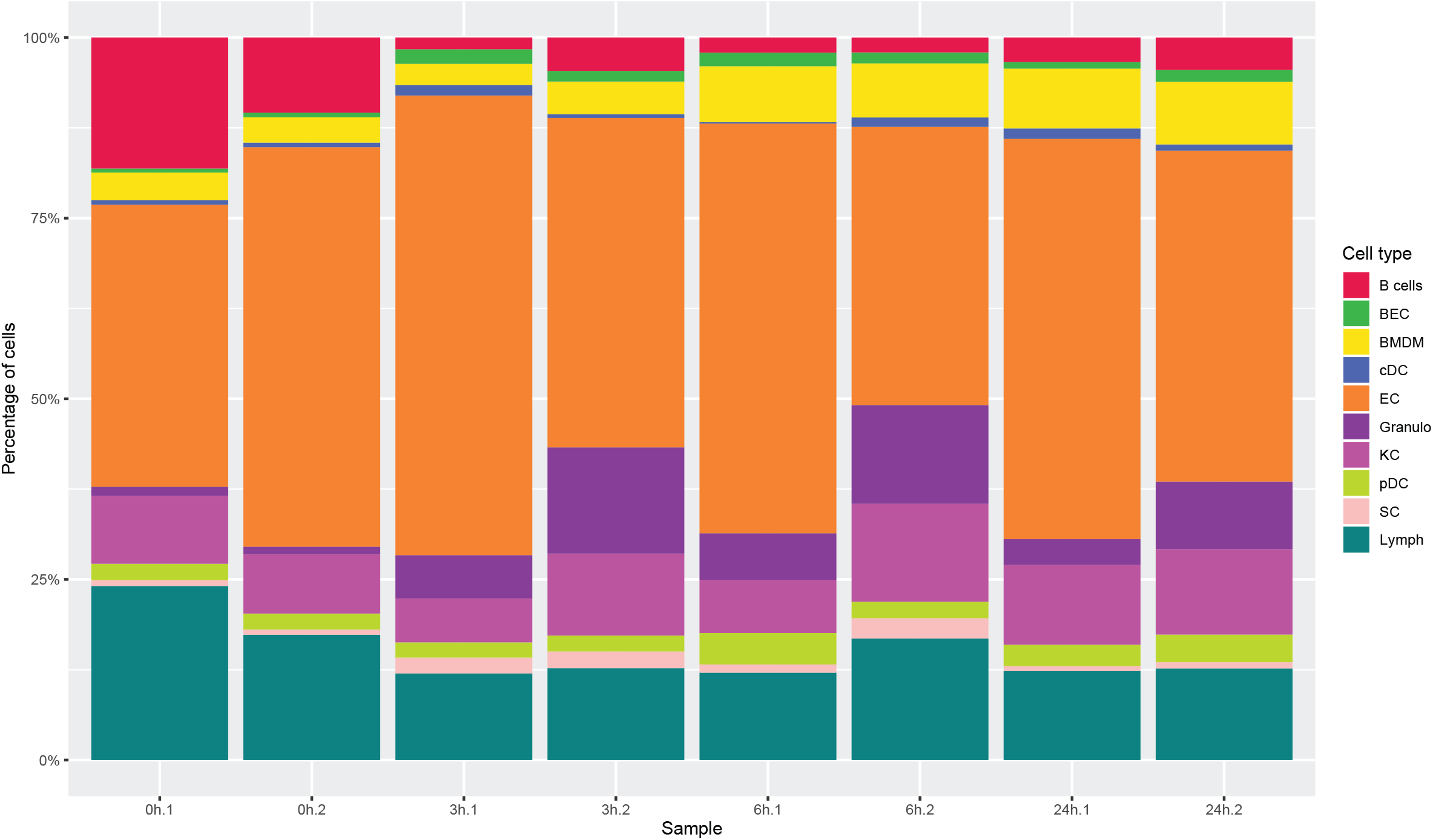
Frequency of cell populations over time. Stack plot showing the percentage of each cell type present in every sample excluding spiked-in hepatocytes (refer to methods).

**Extended Data Fig. 8.**
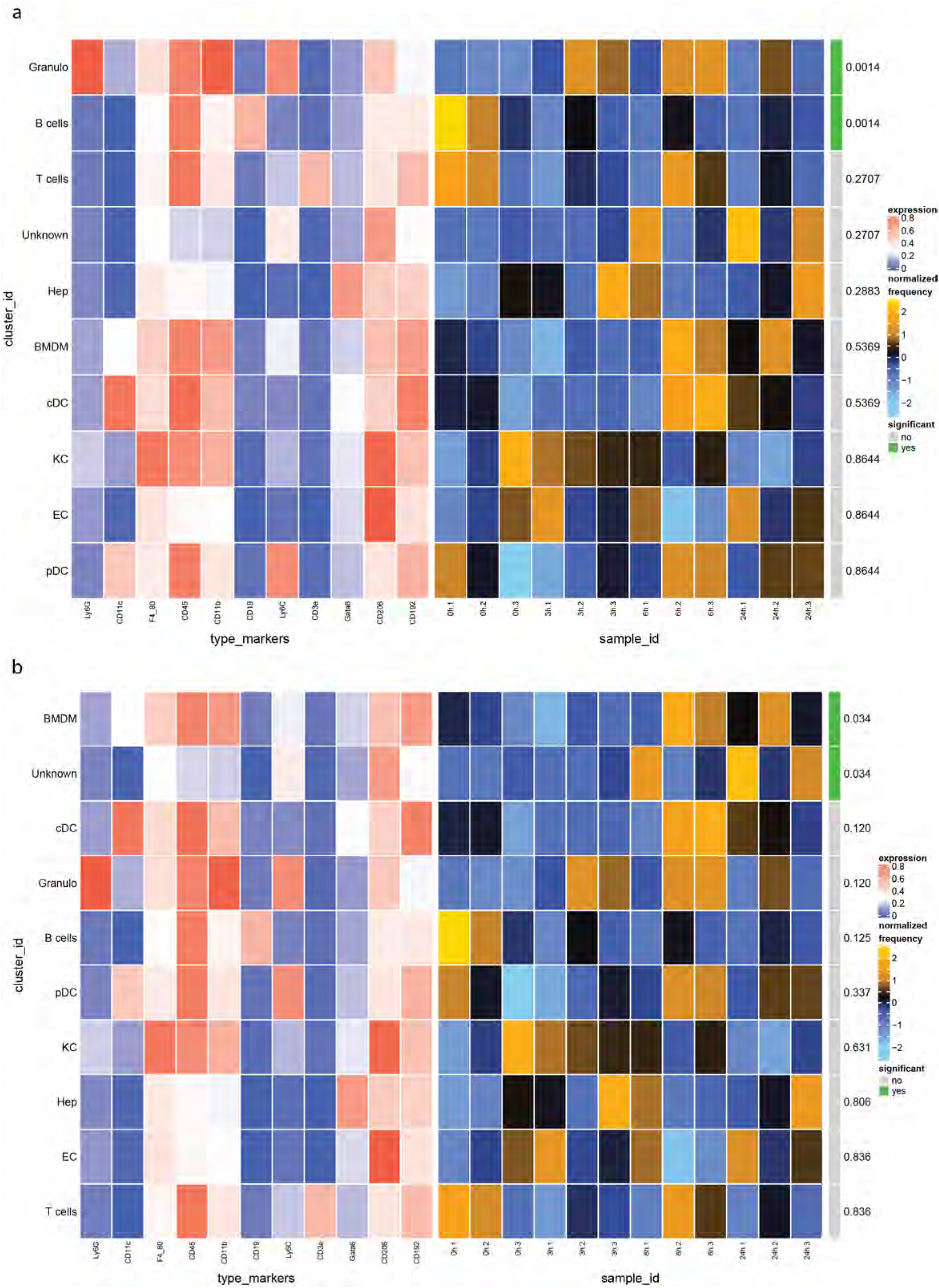
Differential abundance of cells identified by CyTOF. Heatmap of the lineage marker expression and cell frequency of each cell type over time, with statistical significance assessed by diffcyt, a) control compared with 3, 6 and 24 hours, b) control and 3 hours compared with 6 and 24 hours.

**Extended Data Fig. 9.**
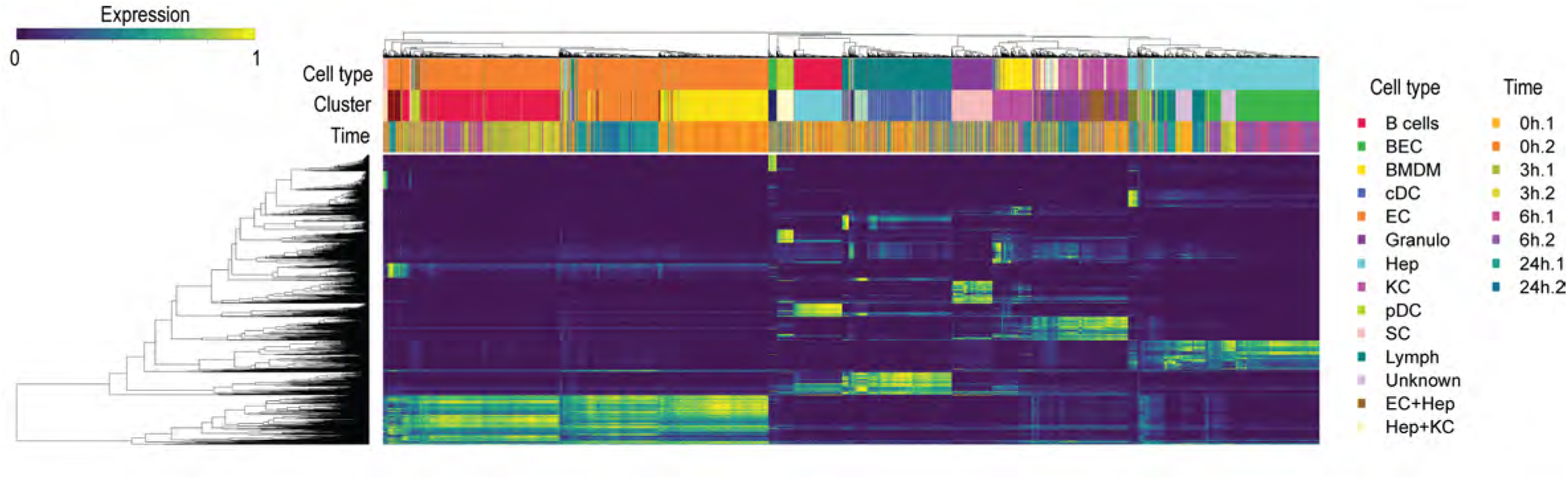
Heatmap gene expression. Heatmap of the MAGIC corrected gene expression of top 50 markers of each unsupervised cluster.

**Extended Data Fig. 10.**
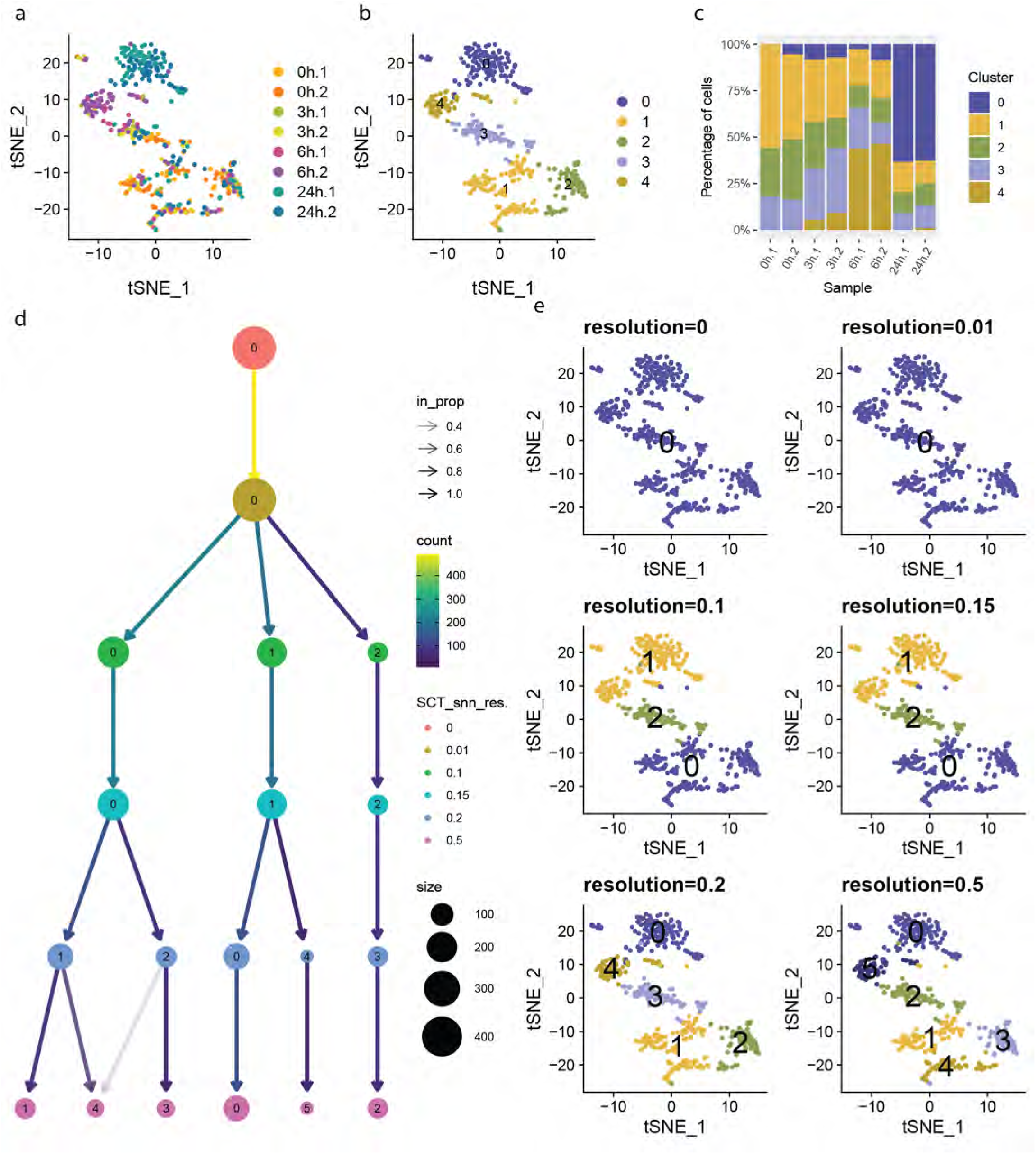
BMDM scRNA-seq data. a) t-SNE visualization of BMDM from scRNA-seq colored by sample, b) by unsupervised cluster. c) stack plot showing the percentage of cells of each unsupervised cluster present in every sample. Cluster 1, 2 and 3 were present at all time points, cluster 4 was mainly present at 6 hours with a small fraction present at 3 hours, cluster 0 was mainly present at 24 hours with a small fraction at all time points. d) Cluster tree with nodes colored by resolution, edges colored by number cells reassigned, node size represents the number of cells in a cluster and transparency is the proportion of cells reassigned. e) t-SNE visualization of BMDM colored by unsupervised clusters with increasing resolution. Firstly, 6 and 24 hours separated from control and 3 hours, secondly 6 and 24 hours separated.

**Extended Data Fig. 11.**
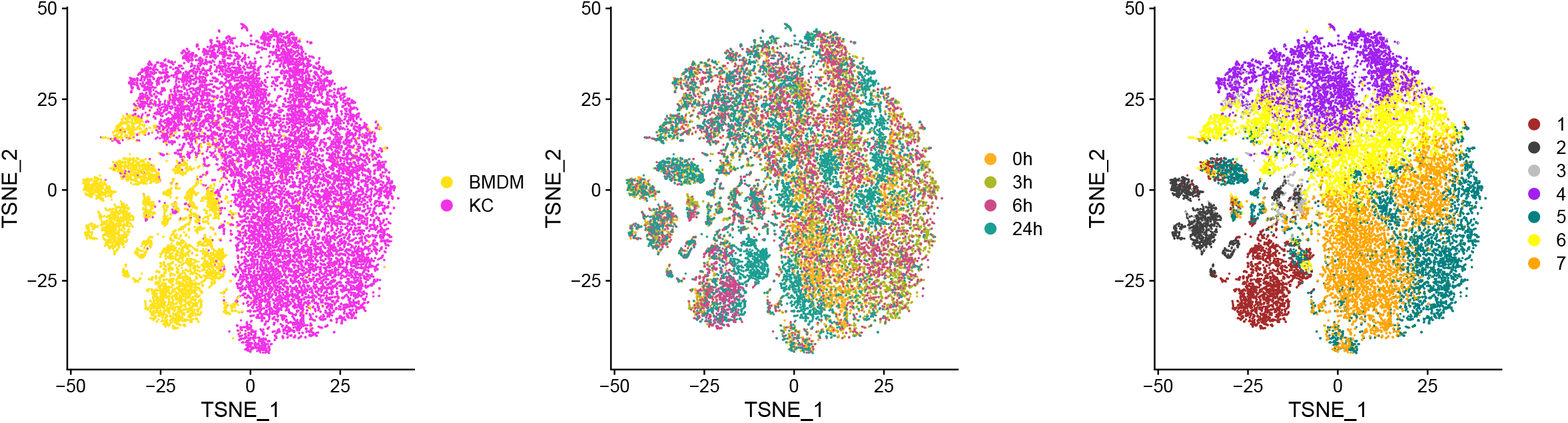
Macrophage CyTOF data. a) t-SNE visualization of Kupffer cells and BMDM from CyTOF colored by cell type, b) by time, c) by unsupervised cluster.

**Extended Data Fig. 12.**
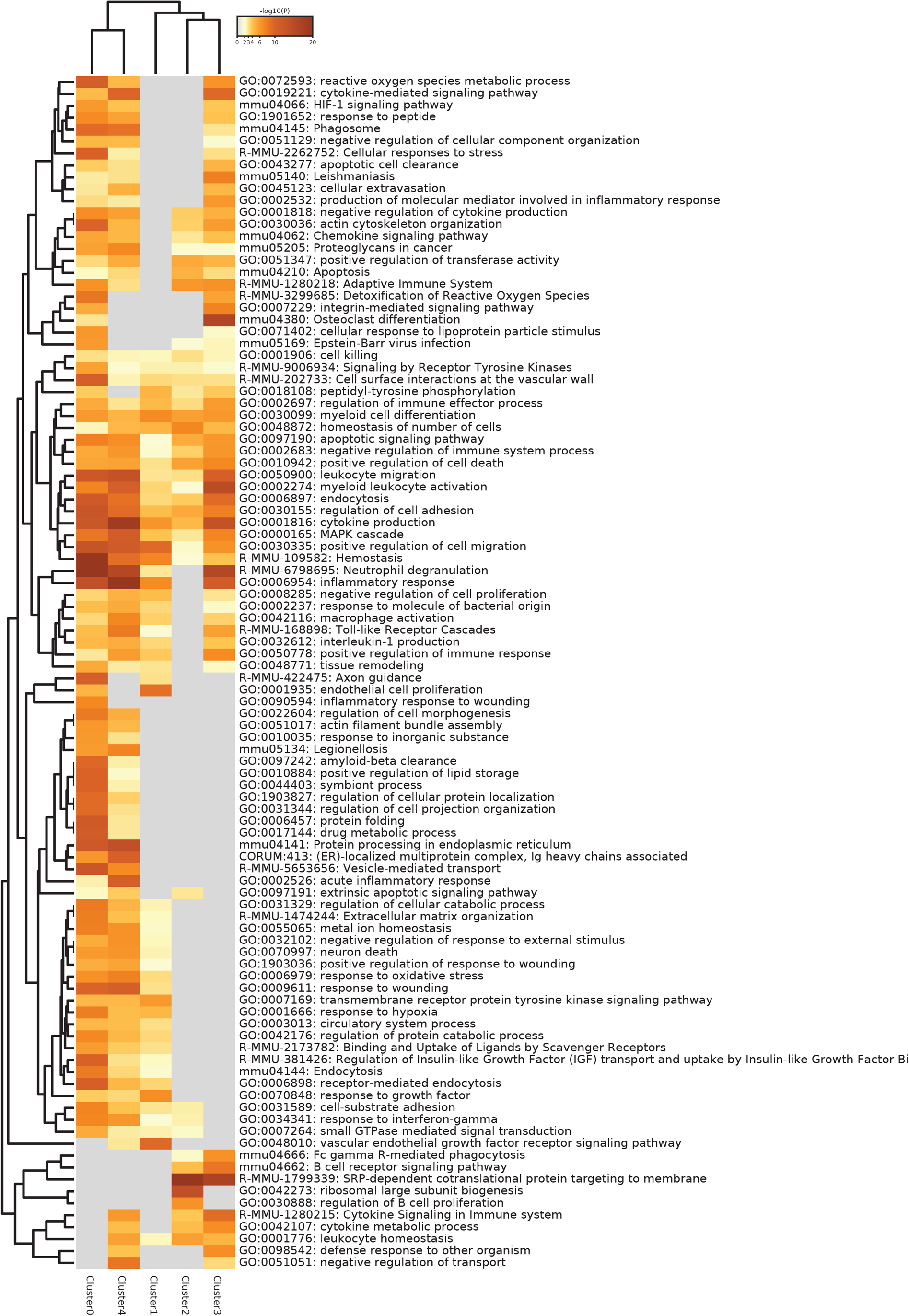
Pathway Enrichment Analysis of BMDM. Top statistically significant pathways associated to the statistically significant genes up regulated in each unsupervised cluster identified by Metascape.

**Extended Data Fig 13.**
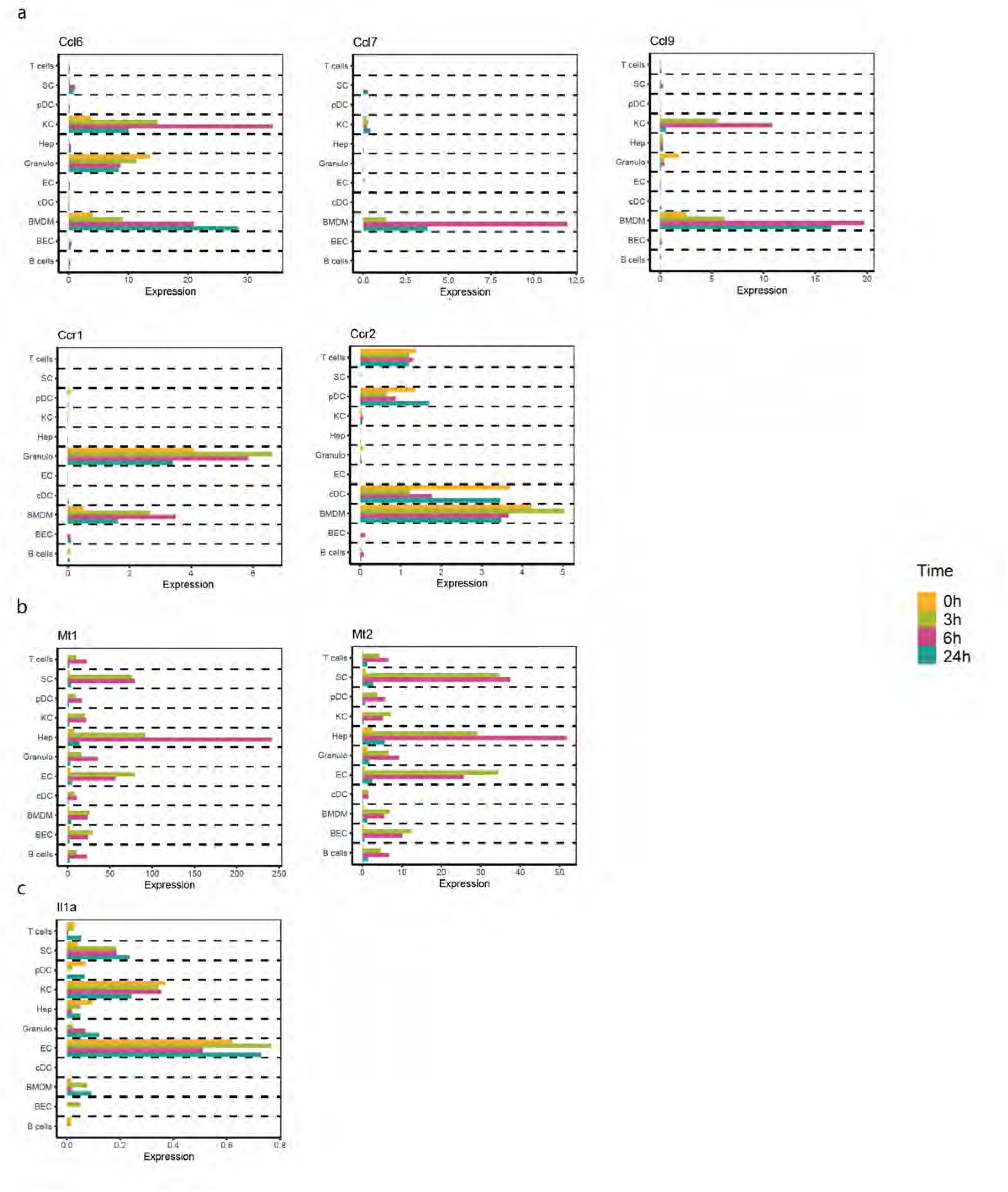
Gene expression of chemokines, cytokines and receptors and antioxidants. Barplot of Ccl6, Ccl7, Ccl9, Ccr1, Ccr2, Mt1, Mt2 and Il1a mRNA.

**Extended Data Fig. 14.**
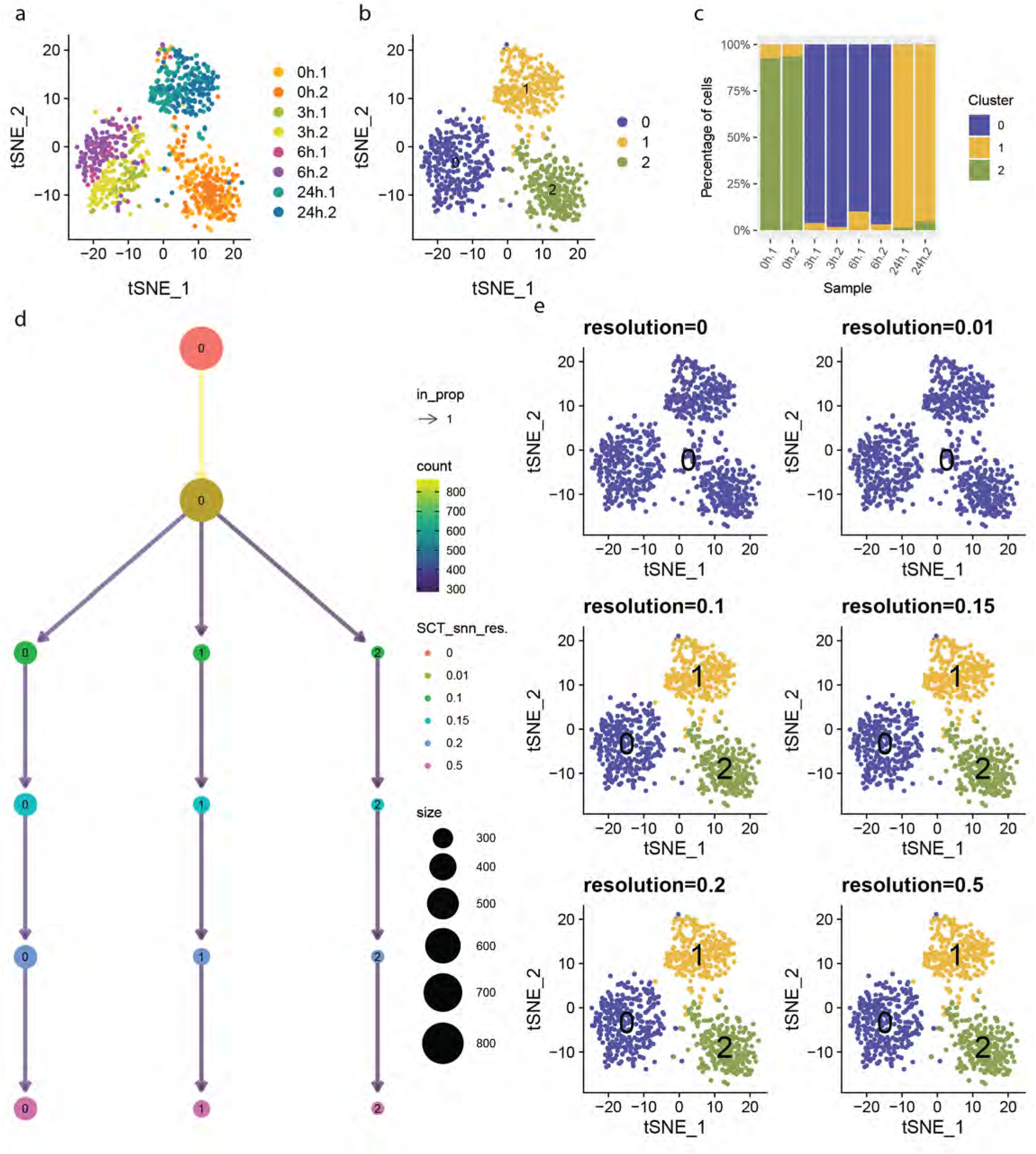
Kupffer cells scRNA-seq data. a) t-SNE visualization of Kupffer cells from scRNA-seq colored by sample, b) by unsupervised cluster. c) stack plot showing the percentage of cells of each unsupervised cluster present in every sample. Cluster 2 was present in 0 hours and a small fraction was present in 24 hours, Cluster 0 was only present at 3 and 6 hours. Cluster 1 appeared at 24 hours post PHx, with a small fraction of cells appearing in all time points. d) Cluster tree with nodes colored by resolution, edges colored by number cells reassigned, node size represents the number of cells in a cluster and transparency is the proportion of cells reassigned. e) t-SNE visualization of Kupffer cells colored by unsupervised clusters with increasing resolution. The three clusters split at a similar resolution without further separation.

**Extended Data Fig. 15.**
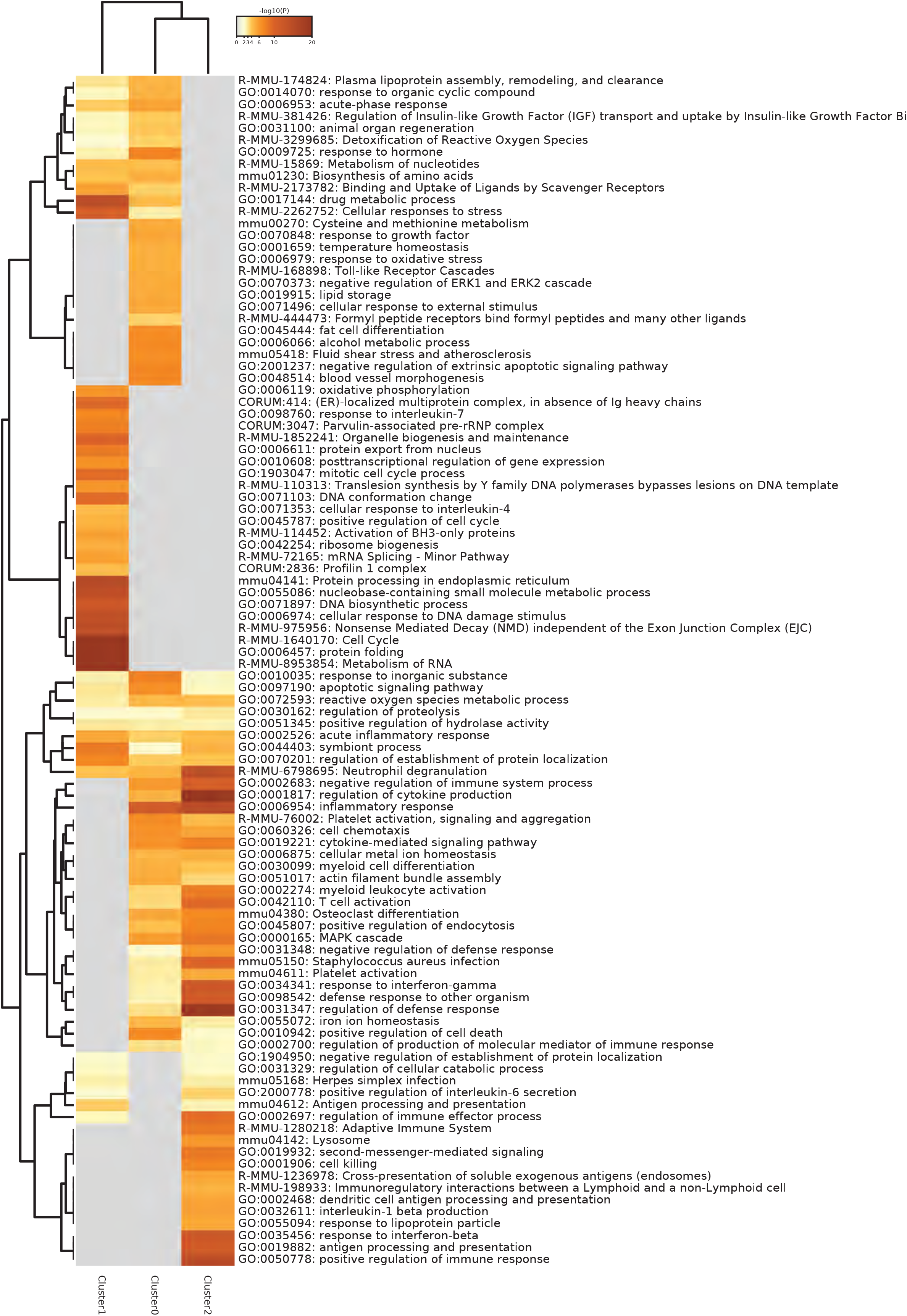
Pathway Enrichment Analysis of Kupffer cells. Top statistically significant pathways associated to the statistically significant genes up regulated in each unsupervised cluster identified by Metascape.

**Extended Data Fig 16.**
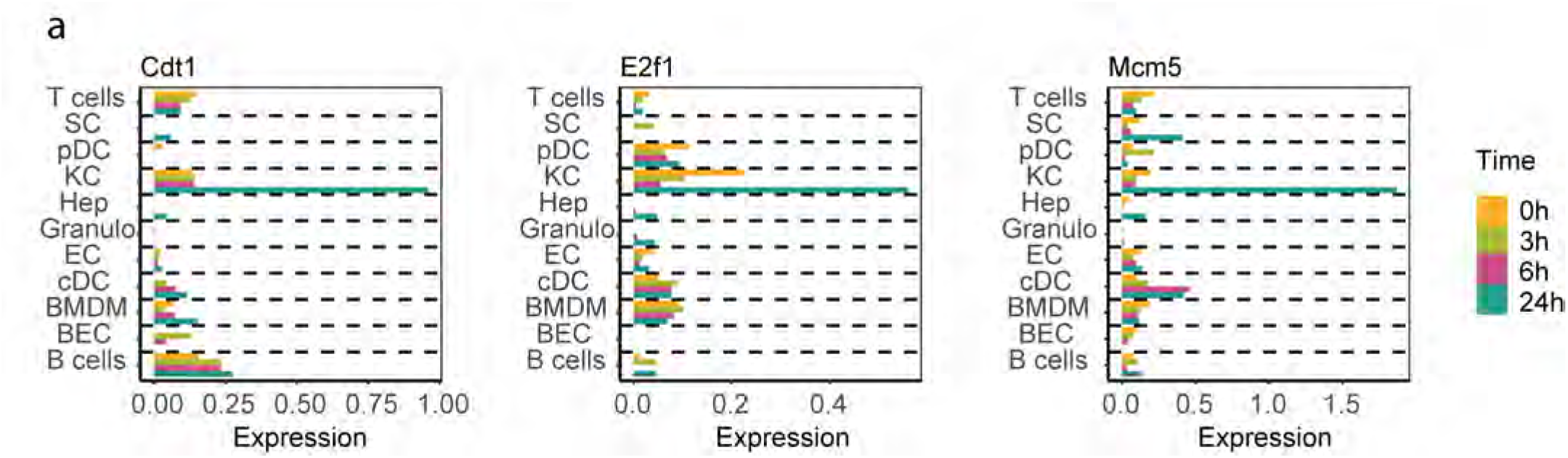
Gene expression of DNA replication genes. Barplots of Cdt1, E2f1, Mcm5.

**Extended Data Fig. 17.**
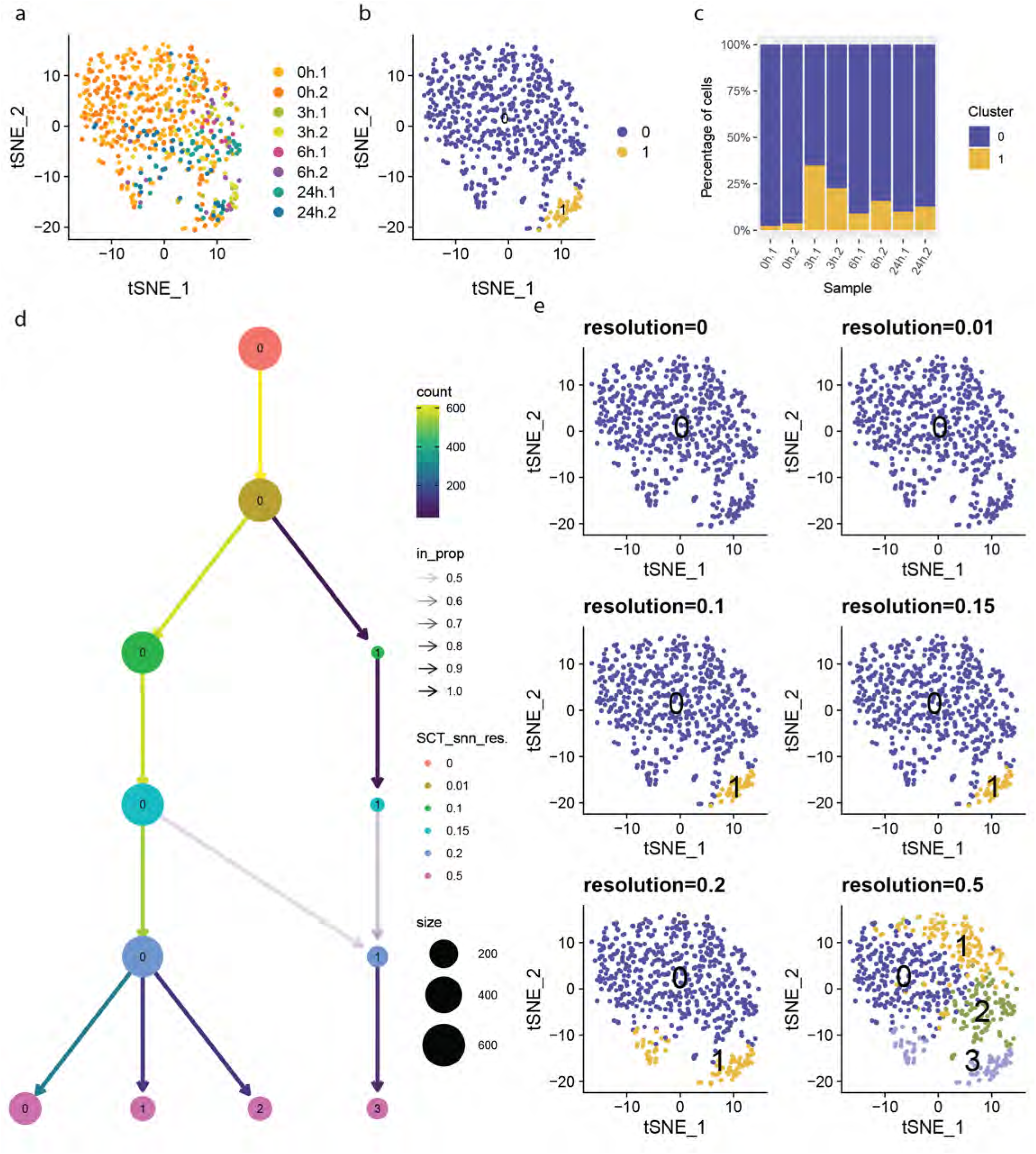
B cell scRNA-seq data. a) t-SNE visualization of B cells from scRNA-seq colored by sample, b) by unsupervised cluster. c) stack plot showing the percentage of cells of each unsupervised cluster present in every sample. d) cluster tree with nodes colored by resolution, edges colored by number cells reassigned, node size represents the number of cells in a cluster and transparency is the proportion of cells reassigned. e) t-SNE visualization of B cells colored by unsupervised clusters with increasing resolution.

**Extended Data Fig. 18.**
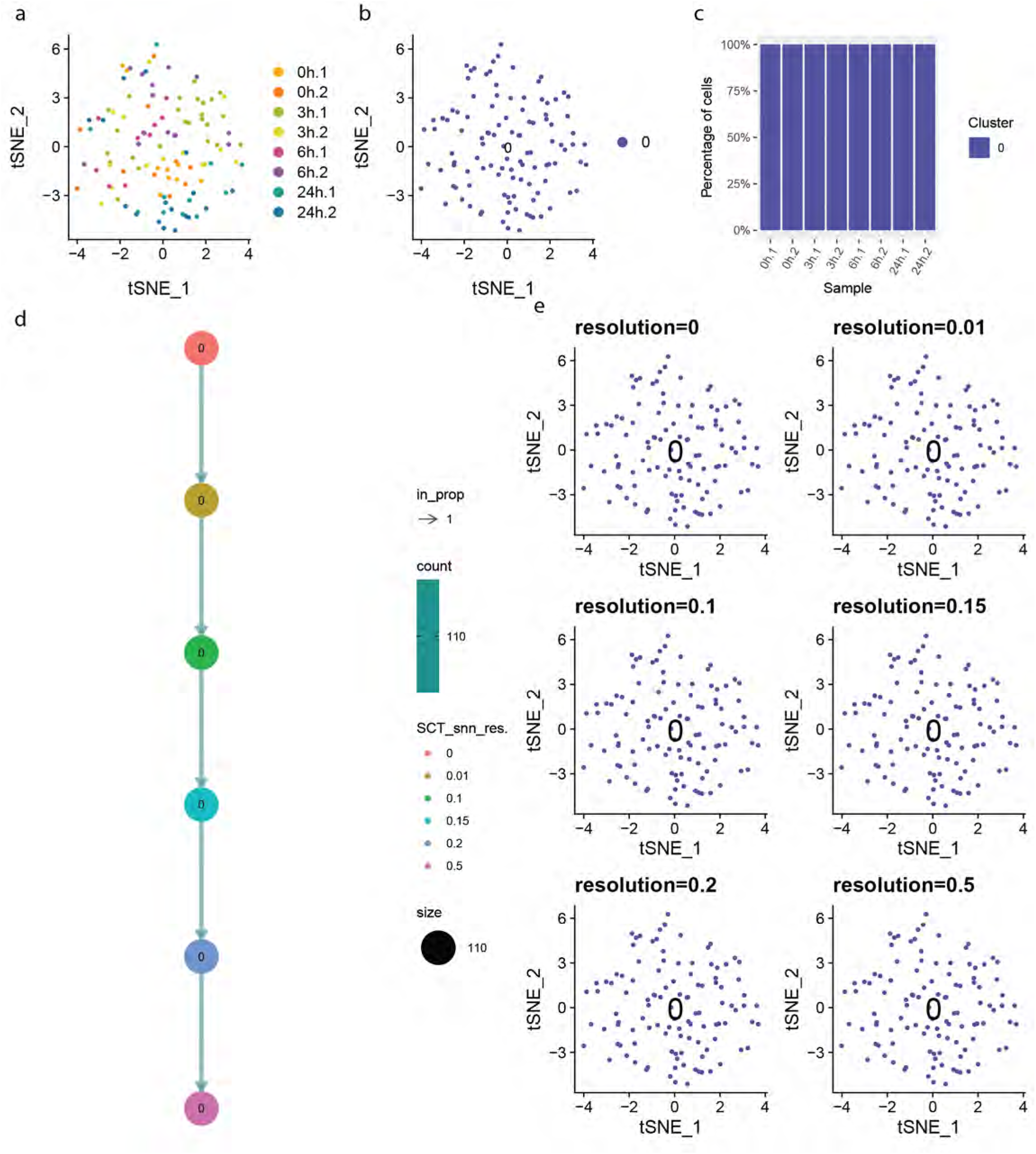
Biliary epithelial cells scRNA-seq data. a) t-SNE visualization of biliary epithelial cells from scRNA-seq colored by sample, b) by unsupervised cluster. c) stack plot showing the percentage of cells of each unsupervised cluster present in every sample. d) cluster tree with nodes colored by resolution, edges colored by number cells reassigned, node size represents the number of cells in a cluster and transparency is the proportion of cells reassigned. e) t-SNE visualization of biliary epithelial cells colored by unsupervised clusters with increasing resolution.

**Extended Data Fig. 19.**
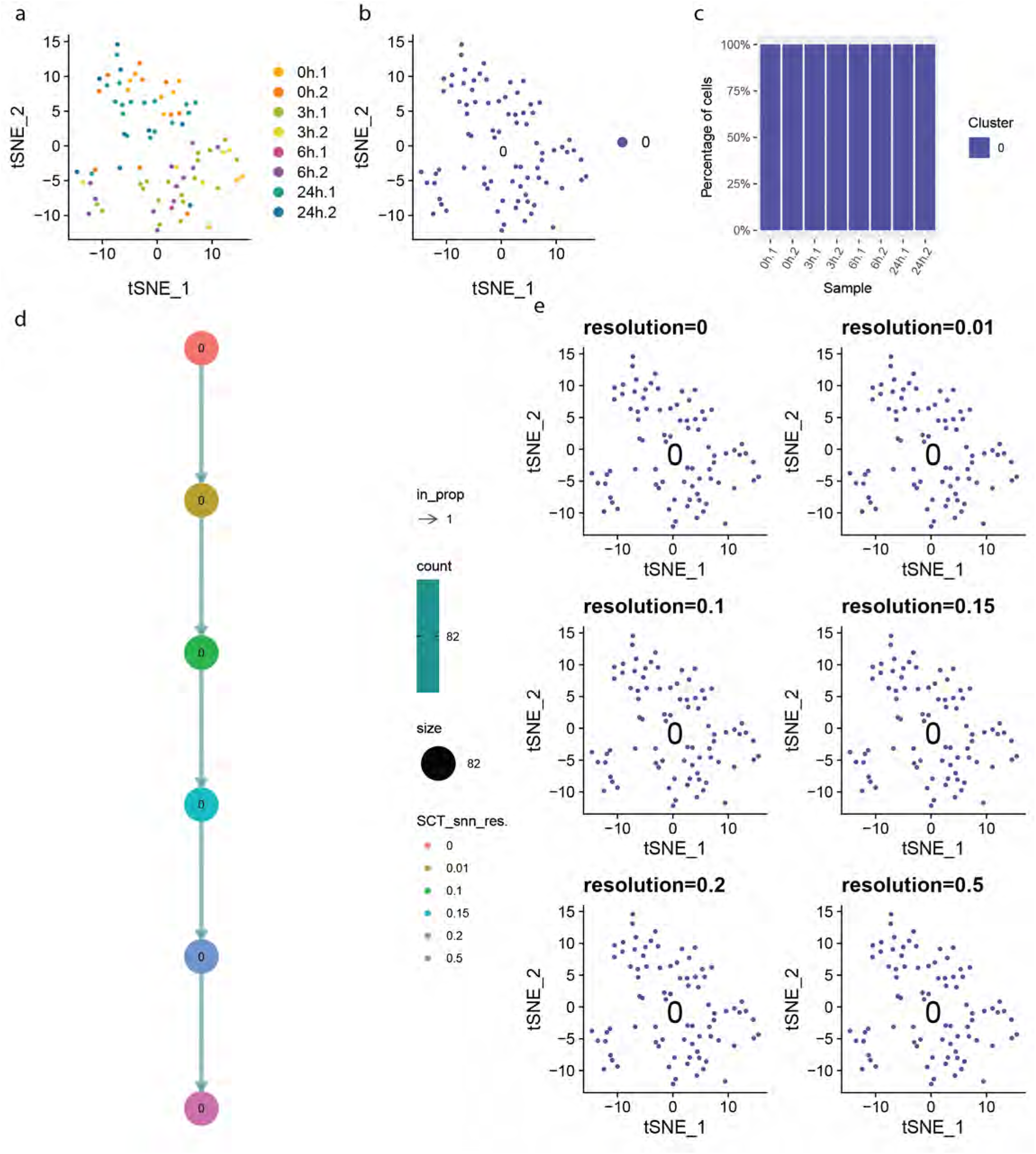
Classic dendritic cells scRNA-seq data. a) t-SNE visualization of classic dendritic cells from scRNA-seq colored by sample, b) by unsupervised cluster. c) stack plot showing the percentage of cells of each unsupervised cluster present in every sample. d) cluster tree with nodes colored by resolution, edges colored by number cells reassigned, node size represents the number of cells in a cluster and transparency is the proportion of cells reassigned. e) t-SNE visualization of classic dendritic cells colored by unsupervised clusters with increasing resolution.

**Extended Data Fig. 20.**
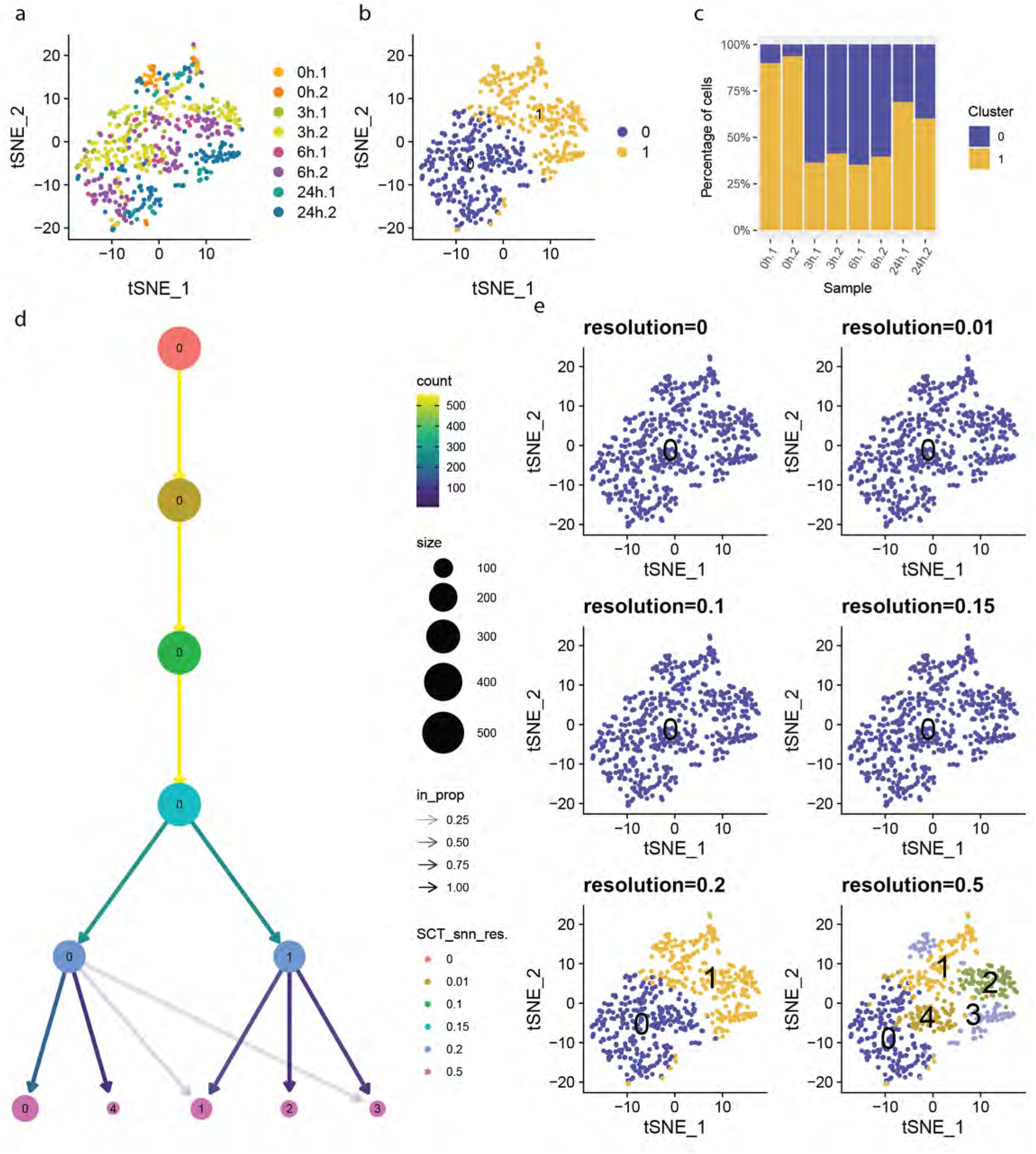
Granulocytes scRNA-seq data. a) t-SNE visualization of granulocytes from scRNA-seq colored by sample, b) by unsupervised cluster. c) stack plot showing the percentage of cells of each unsupervised cluster present in every sample. d) cluster tree with nodes colored by resolution, edges colored by number cells reassigned, node size represents the number of cells in a cluster and transparency is the proportion of cells reassigned. e) t-SNE visualization of granulocytes colored by unsupervised clusters with increasing resolution.

**Extended Data Fig. 21.**
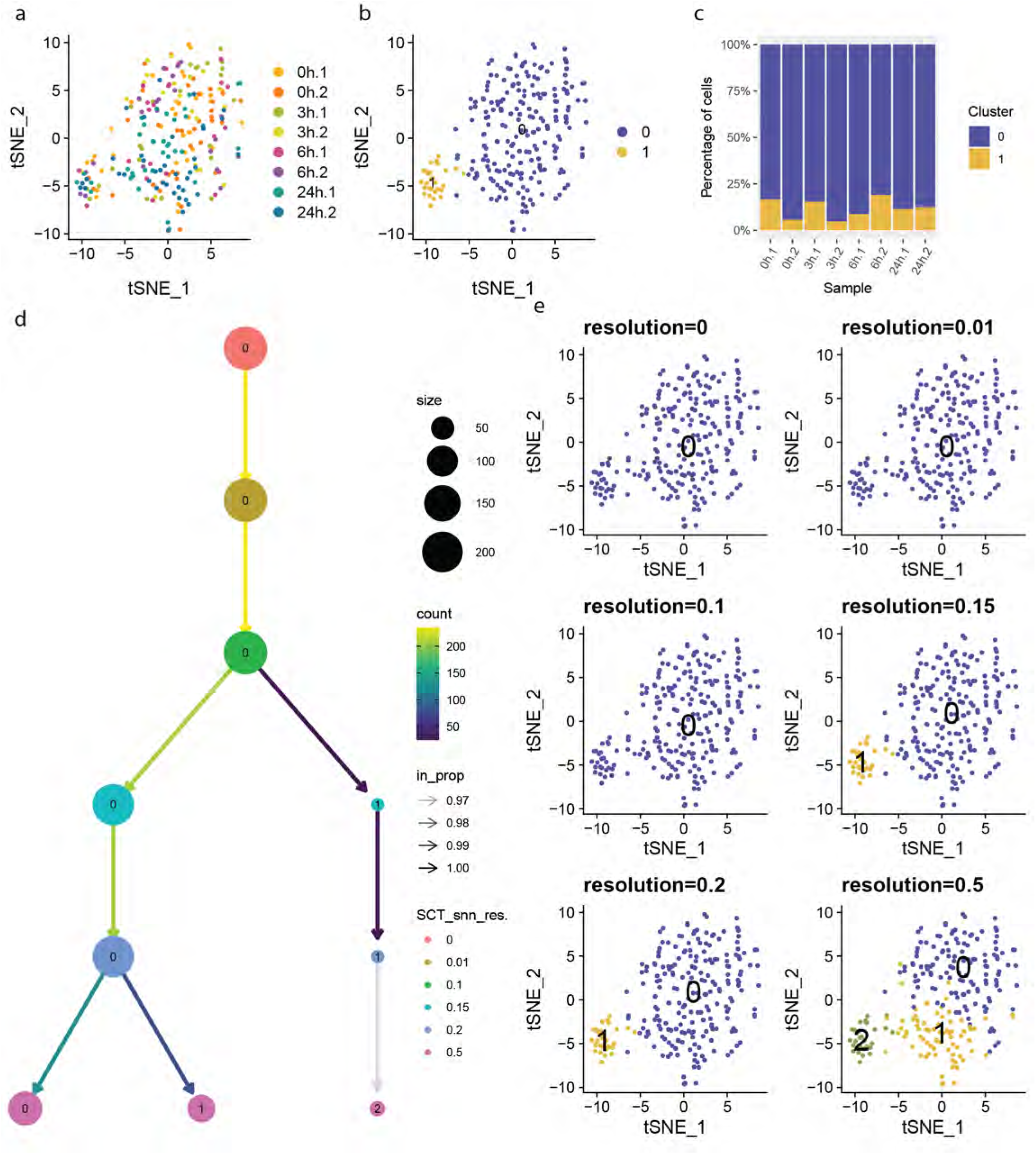
Plasmacytoid dendritic cells scRNA-seq data. a) t-SNE visualization of pDC from scRNA-seq colored by sample, b) by unsupervised cluster. c) stack plot showing the percentage of cells of each unsupervised cluster present in every sample. d) cluster tree with nodes colored by resolution, edges colored by number cells reassigned, node size represents the number of cells in a cluster and transparency is the proportion of cells reassigned. e) t-SNE visualization of plasmacytoid dendritic cells colored by unsupervised clusters with increasing resolution.

**Extended Data Fig. 22.**
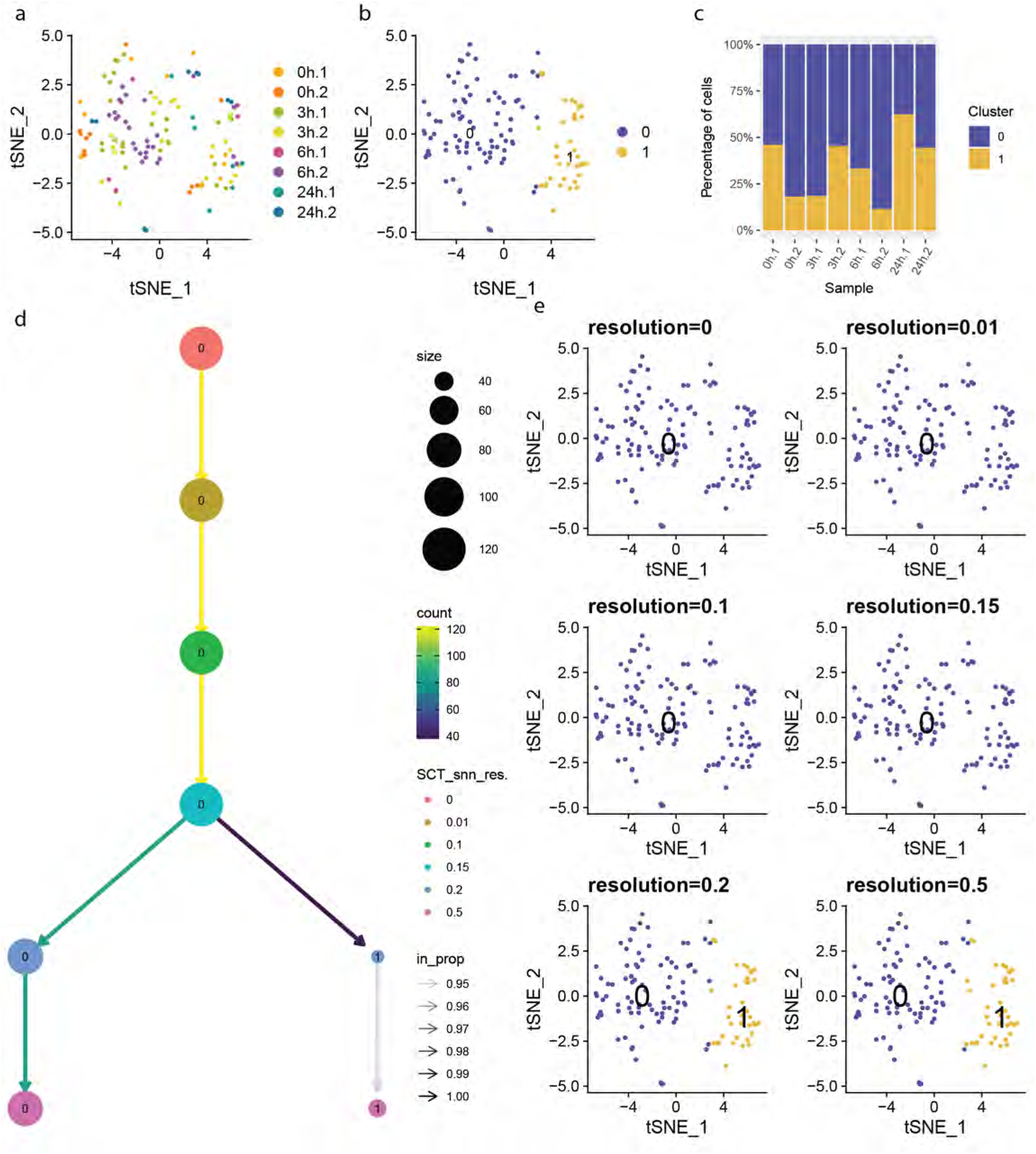
Stellate cell scRNA-seq data. a) t-SNE visualization of stellate cell from scRNA-seq colored by sample, b) by unsupervised cluster. c) stack plot showing the percentage of cells of each unsupervised cluster present in every sample. d) cluster tree with nodes colored by resolution, edges colored by number cells reassigned, node size represents the number of cells in a cluster and transparency is the proportion of cells reassigned. e) t-SNE visualization of stellate cell colored by unsupervised clusters with increasing resolution.

**Extended Data Fig. 23.**
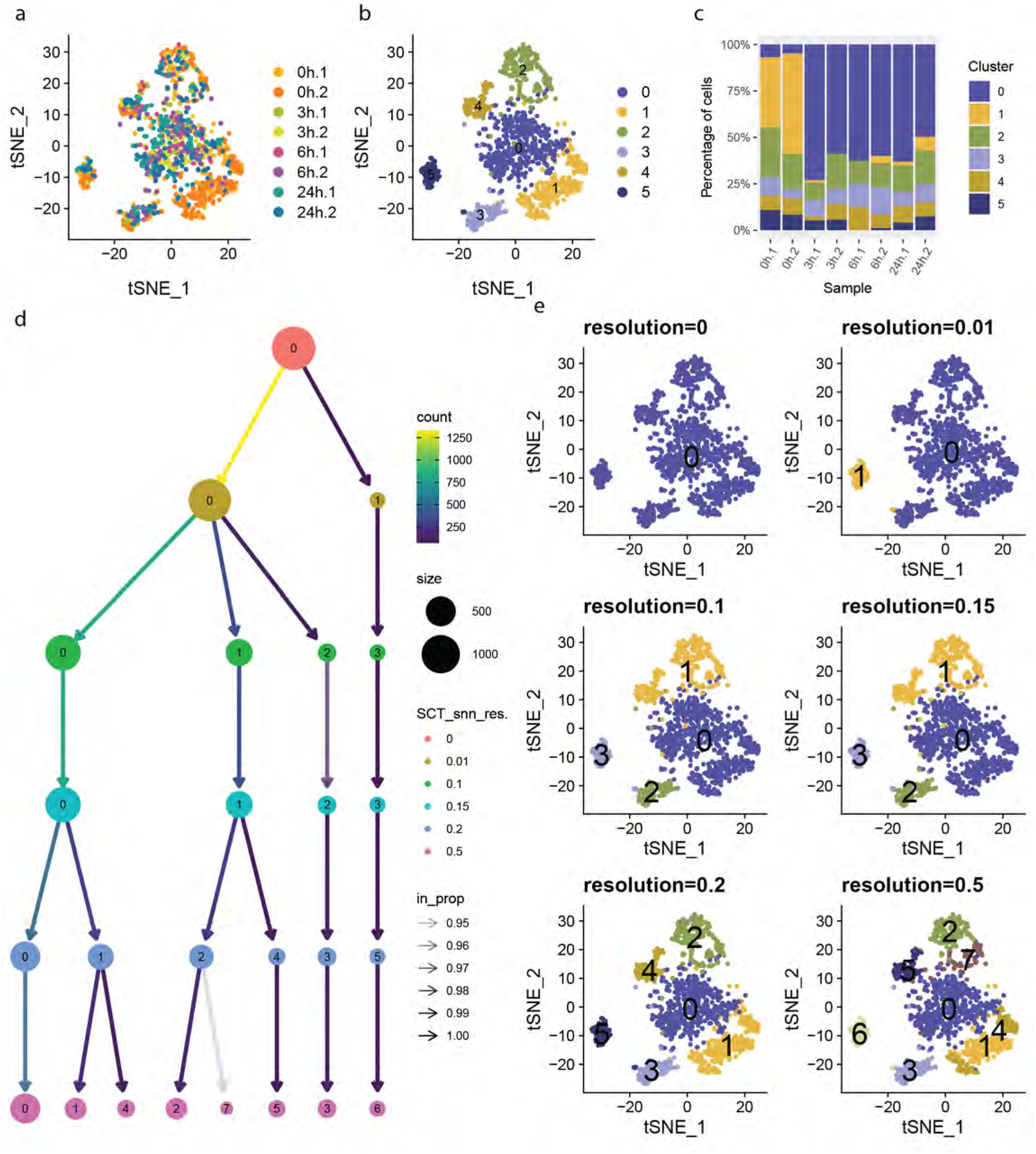
Lymphocytes scRNA-seq data. a) t-SNE visualization of T cells from scRNA-seq colored by sample, b) by unsupervised cluster. c) stack plot showing the percentage of cells of each unsupervised cluster present in every sample. d) cluster tree with nodes colored by resolution, edges colored by number cells reassigned, node size represents the number of cells in a cluster and transparency is the proportion of cells reassigned. e) t-SNE visualization of T cells colored by unsupervised clusters with increasing resolution.

**Extended Data Fig. 24.**
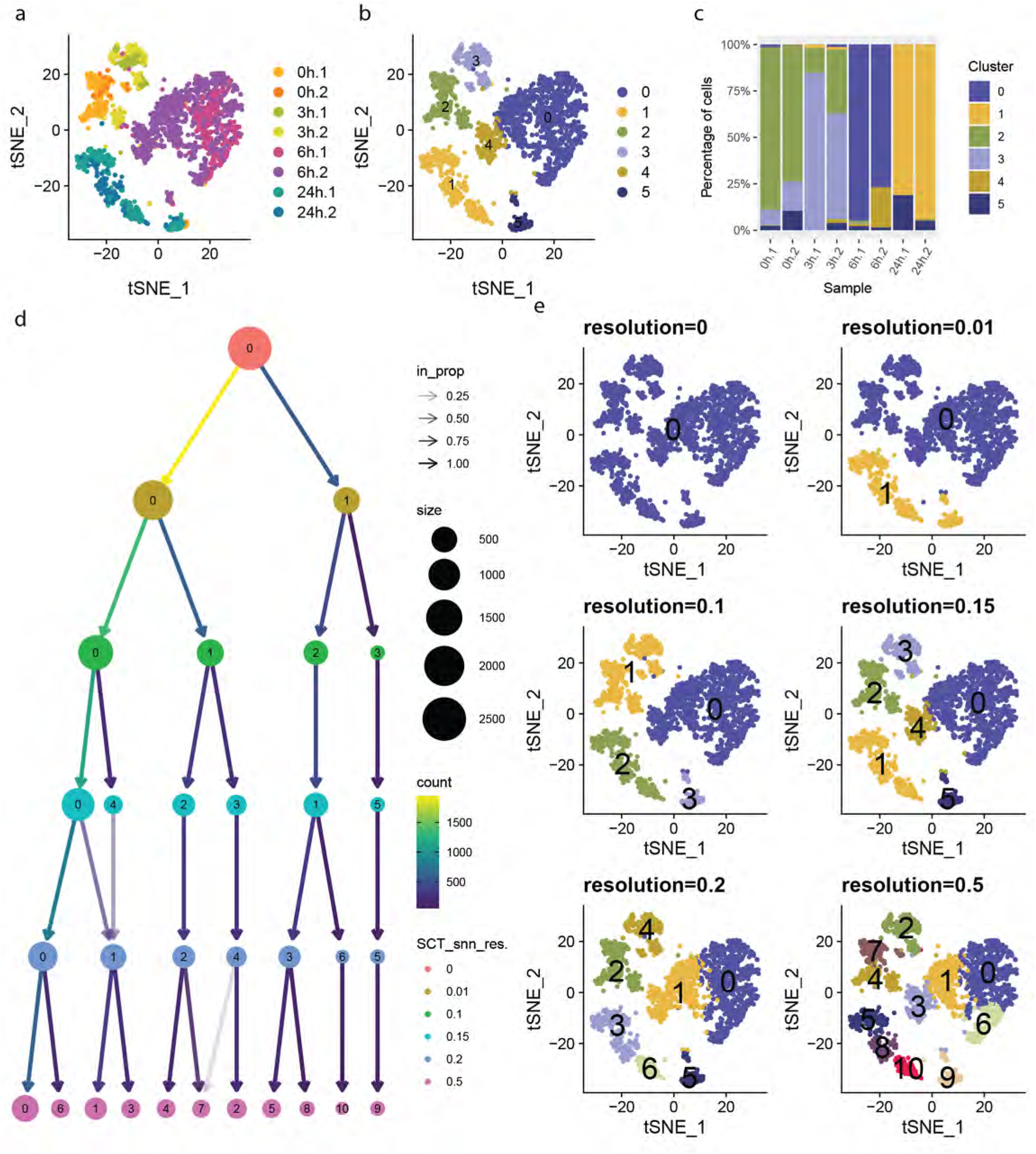
Hepatocyte scRNA-seq data. a) t-SNE visualization of hepatocytes from scRNA-seq colored by sample, b) by unsupervised cluster. c) stack plot showing the percentage of cells of each unsupervised cluster present in every sample. Cluster 2 was present mainly in control samples and a small fraction in 3 hours, Cluster 3 mainly present at 3 hours and a small fraction in control. Clusters 0 and 4 are present at 6 hours, and Cluster 1 appears at 24 hours. Finally, Cluster 5 appears as a small fraction present in each time point. d) Cluster tree with nodes colored by resolution, edges colored by number cells reassigned, node size represents the number of cells in a cluster and transparency is the proportion of cells reassigned. e) t-SNE visualization of hepatocytes colored by unsupervised clusters with increasing resolution. 24 hours first separated from the others, secondly 6 hours separated from control and 3 hours, and finally, 3 hours and the control separated.

**Extended Data Fig. 25.**
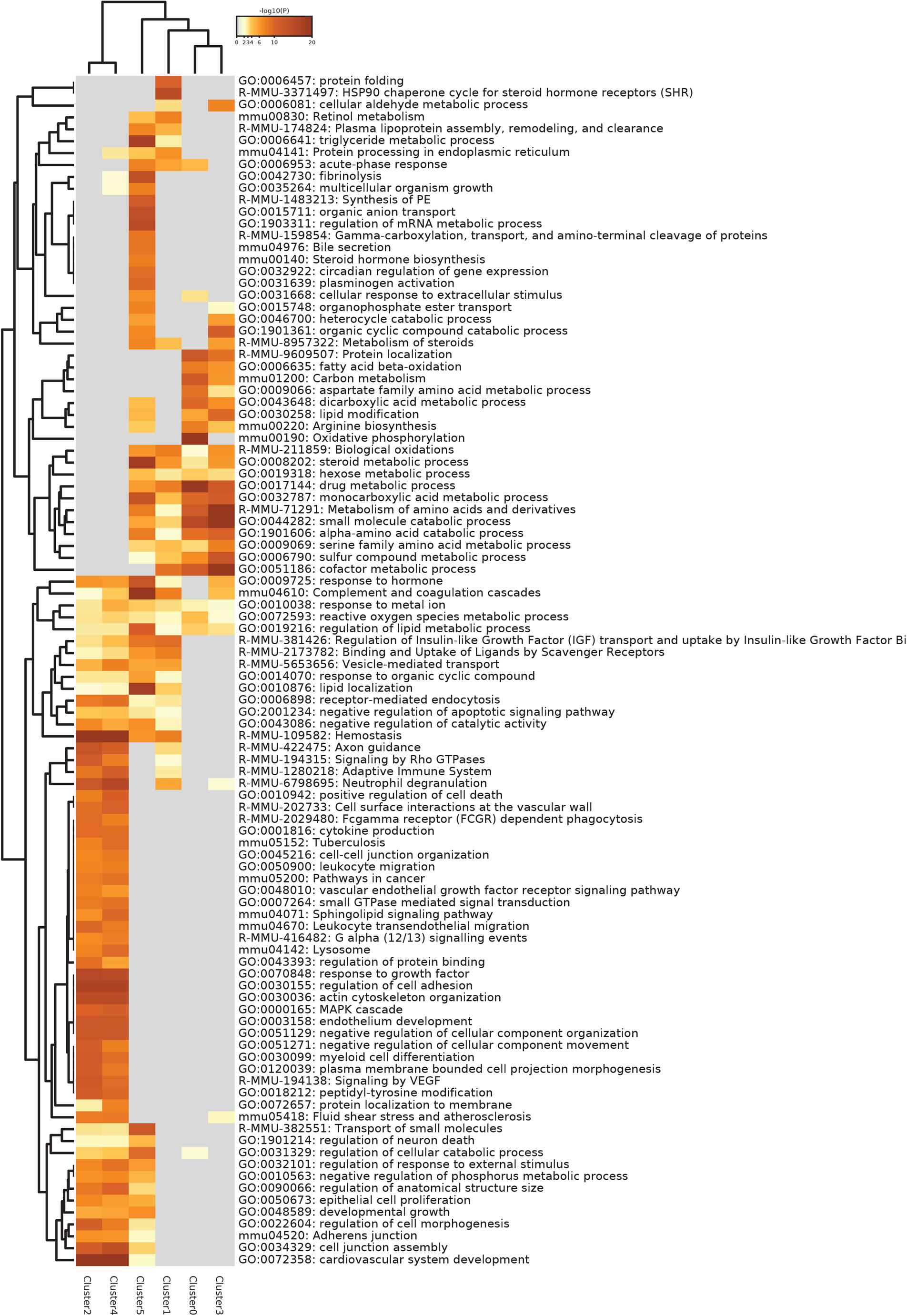
Pathway Enrichment Analysis of Hep. Top statistically significant pathways associated to the statistically significant genes up regulated in each unsupervised cluster identified by Metascape.

**Extended Data Fig. 26.**
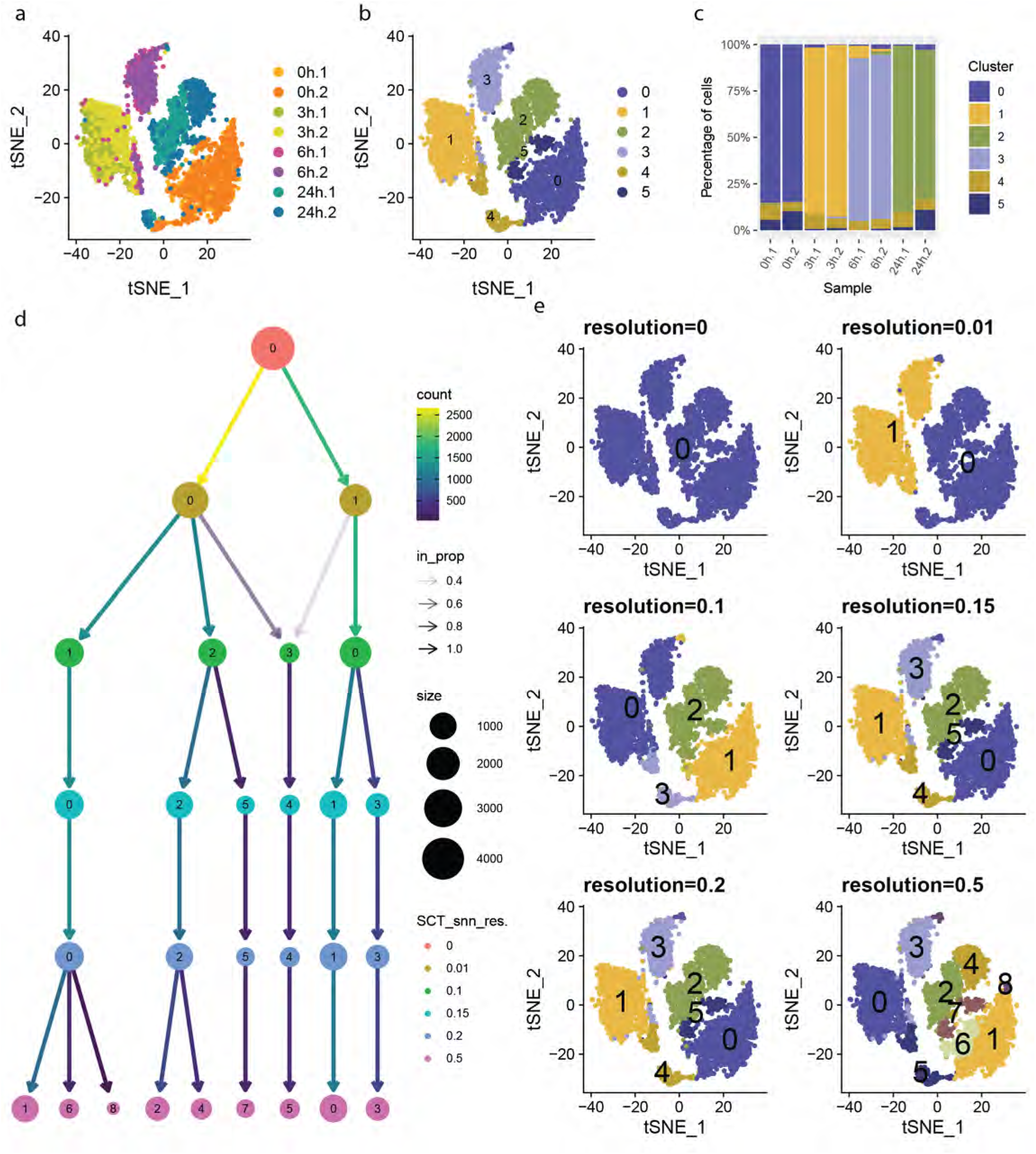
Endothelial cell scRNA-seq data. a) t-SNE visualization of endothelial cells from scRNA-seq colored by sample, b) by unsupervised cluster. c) stack plot showing the percentage of cells of each unsupervised cluster present in every sample. Cluster 0 was only present in the control samples, Cluster 1 coincided with 3 hours post PHx, Cluster 3 with 6 hours post PHx, Cluster 2 appeared at 24 hours post PHx. Finally, Cluster 4 and 5 appeared as a small fraction present in each time point. d) cluster tree with nodes colored by resolution, edges colored by number cells reassigned, node size represents the number of cells in a cluster and transparency is the proportion of cells reassigned. e) t-SNE visualization of endothelial cells colored by unsupervised clusters with increasing resolution. Control and 24 hours separated from 3 and 6 hours, secondly 24 hour separated from control and finally, 3 hours separated from 6 hours.

**Extended Data Fig. 27.**
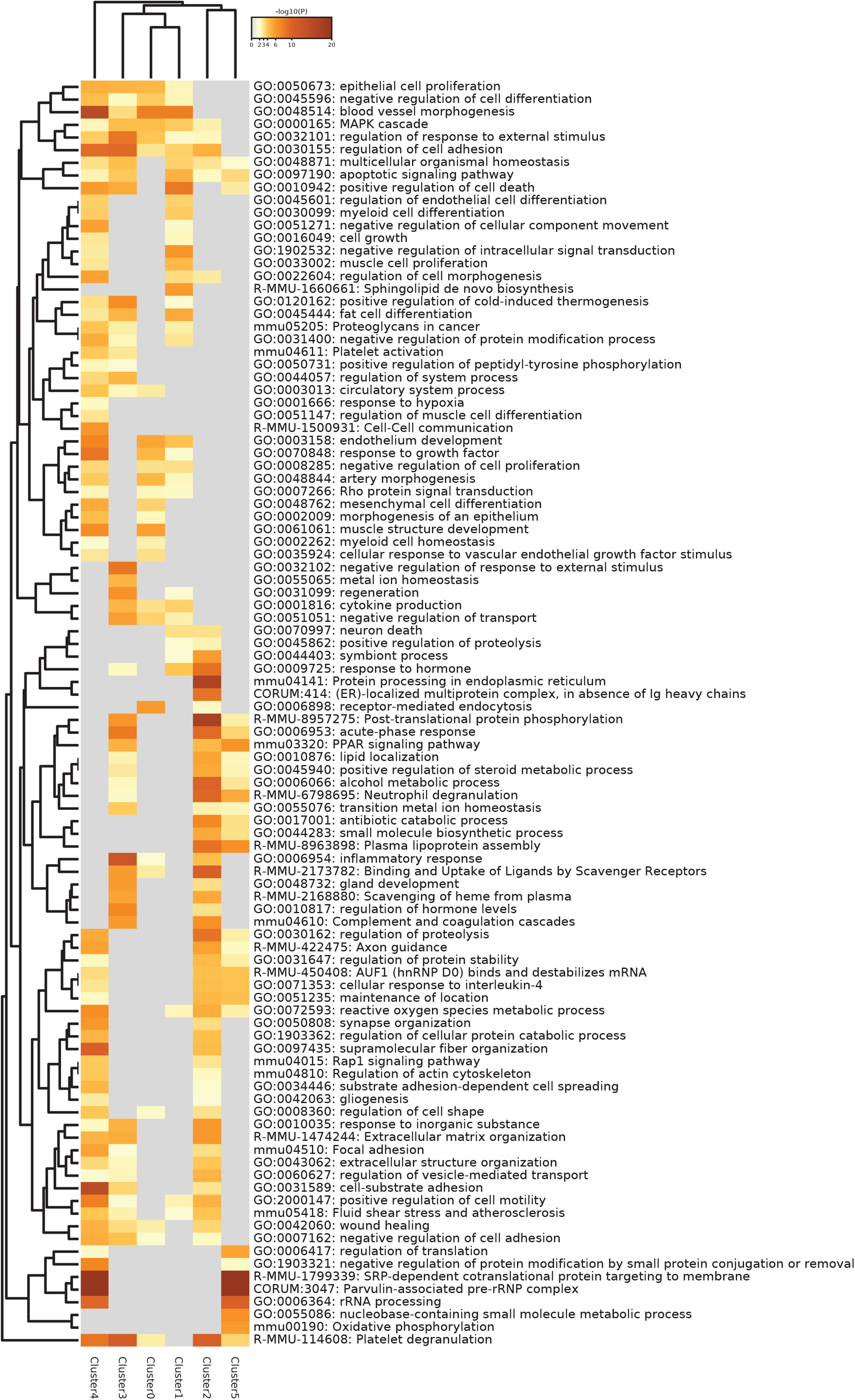
EC Pathway Enrichment Analysis of EC. Top statistically significant pathways associated to the statistically significant genes up regulated in each unsupervised cluster identified by Metascape.

**Extended Data Fig. 28.**
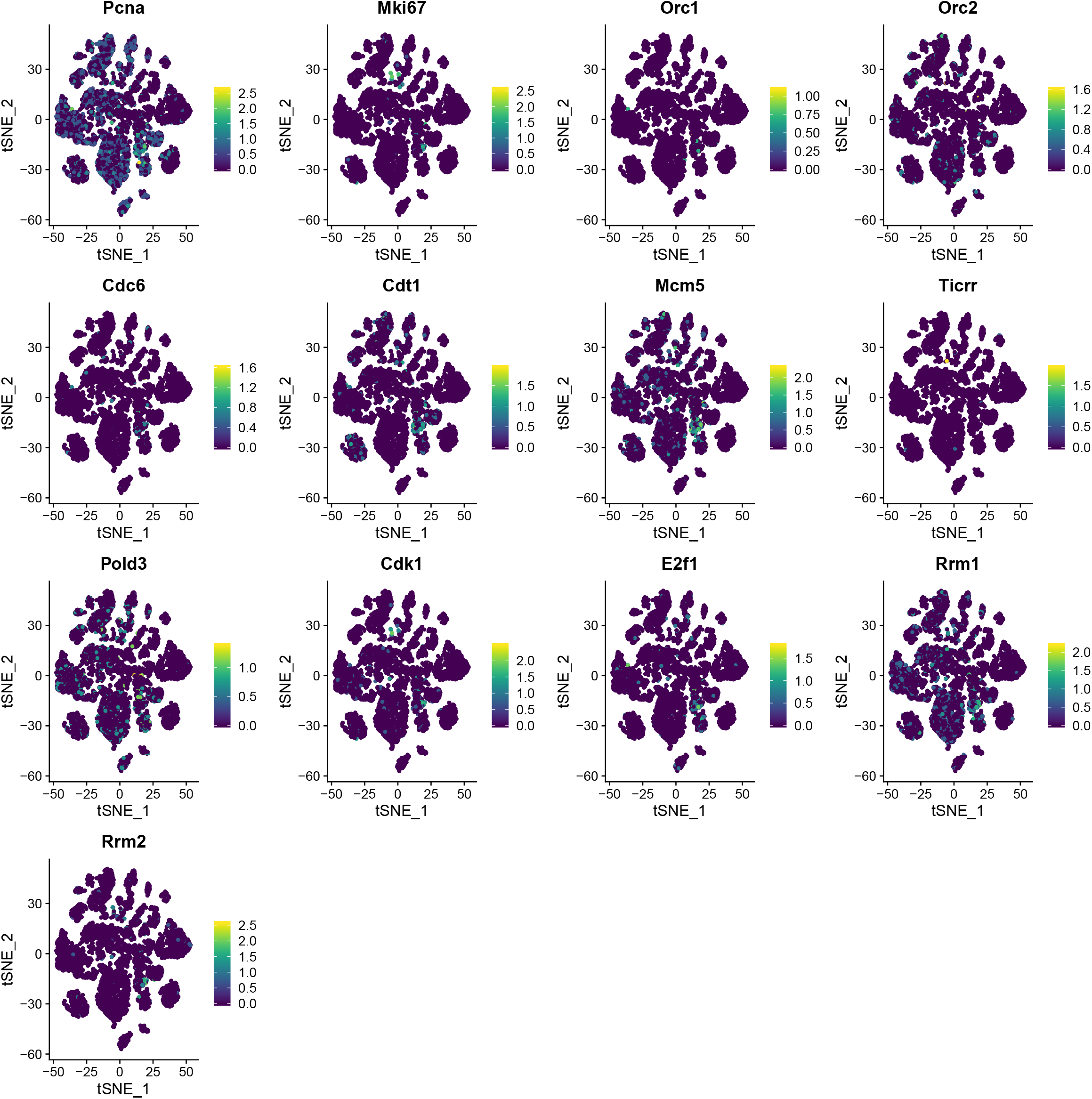
DNA replication genes. t-SNE visualization of the gene expression of the DNA replication genes.

**Extended Data Fig. 29.**
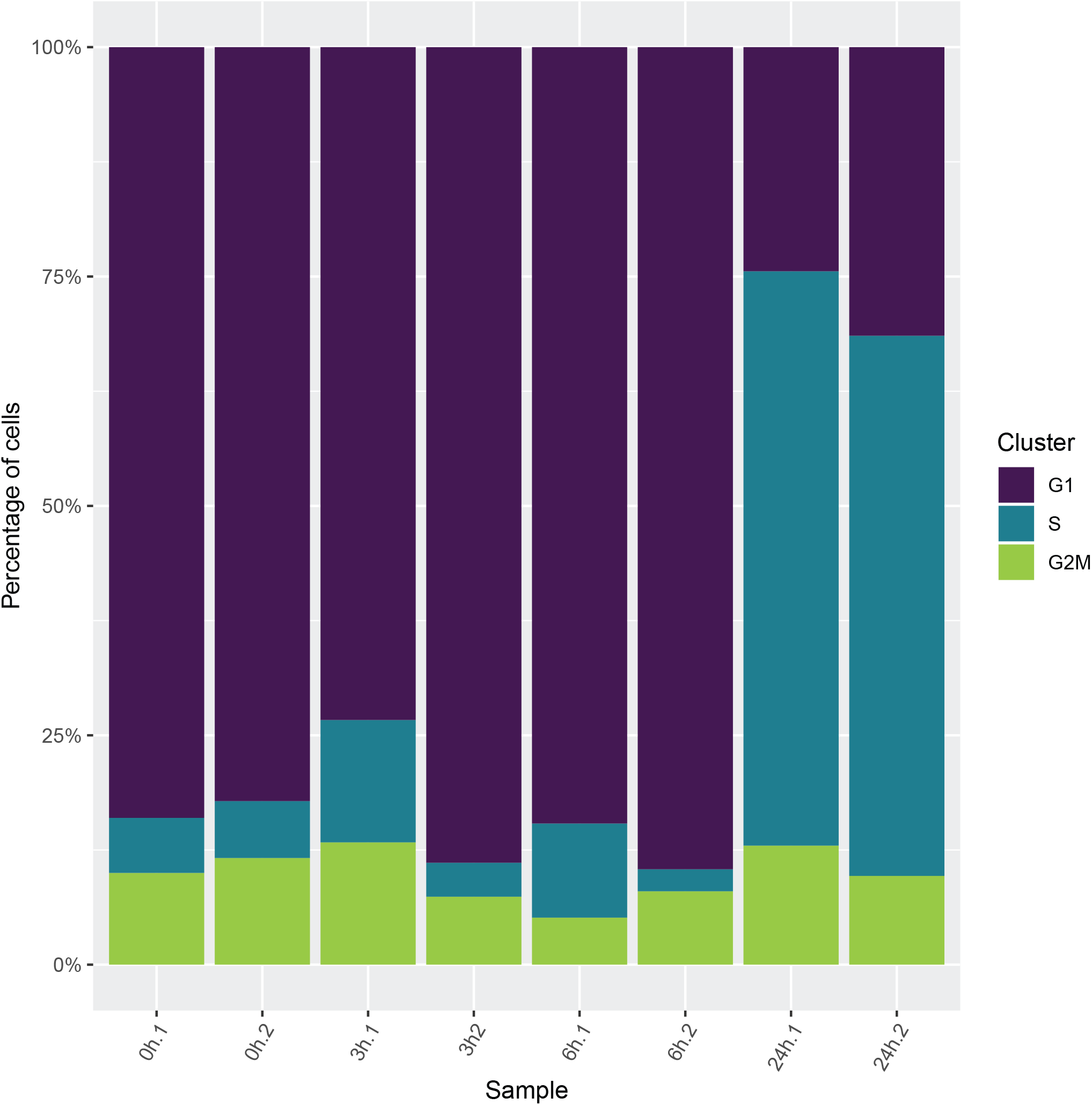
Cell cycle genes in Kupffer cells. Stack plot showing the percentage of Kupffer cells assigned to each cell cycle phase.

**Extended Data Fig. 30:**
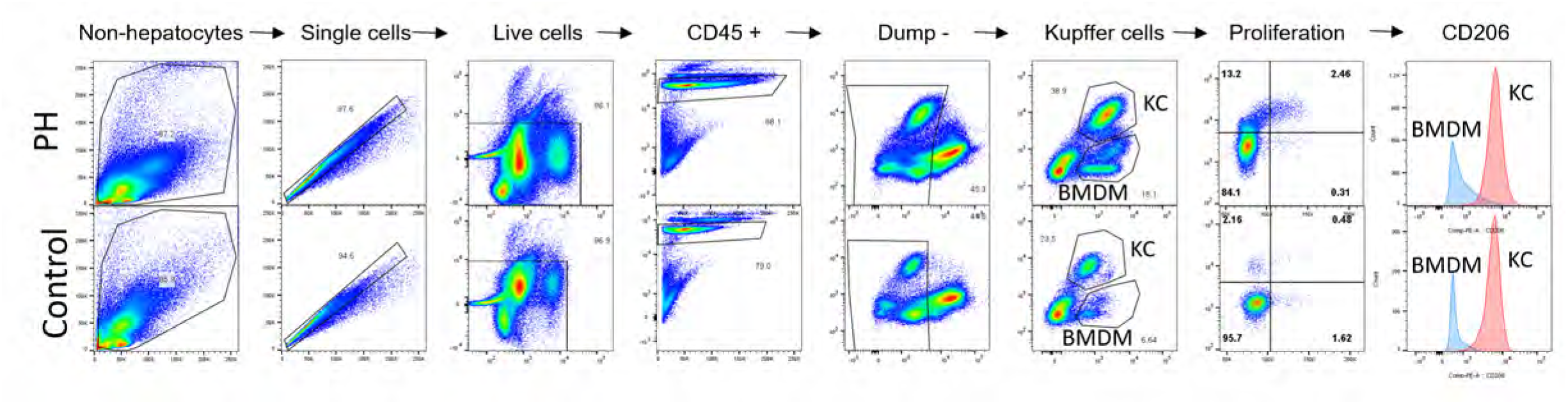
Gating strategy for DNA integration of EdU. Single cells were obtained from murine livers in control (no PHx) and 26 hour post-PHx. All non-hepatocytes were gated, then single cells and live cells. CD45 positive were selected and B cells (CD19), T cells (CD3), endothelial cells (CD31), and NK cells (NK1.1) were gated out with the dump channel. F4/80 positive CD11b positive cells were identified as Kupffer cells and proliferation was measured with EdU intercalation and DNA (FxCycle Violet) content in this population. CD206 expression in Kupffer cells (red) overlaid in a histogram against remaining BMDM CD11b+/F4/80 dim/-populations (blue) confirms Kupffer cells are CD206+ and BMDM are CD206-.

**Extended Data Fig. 31.**
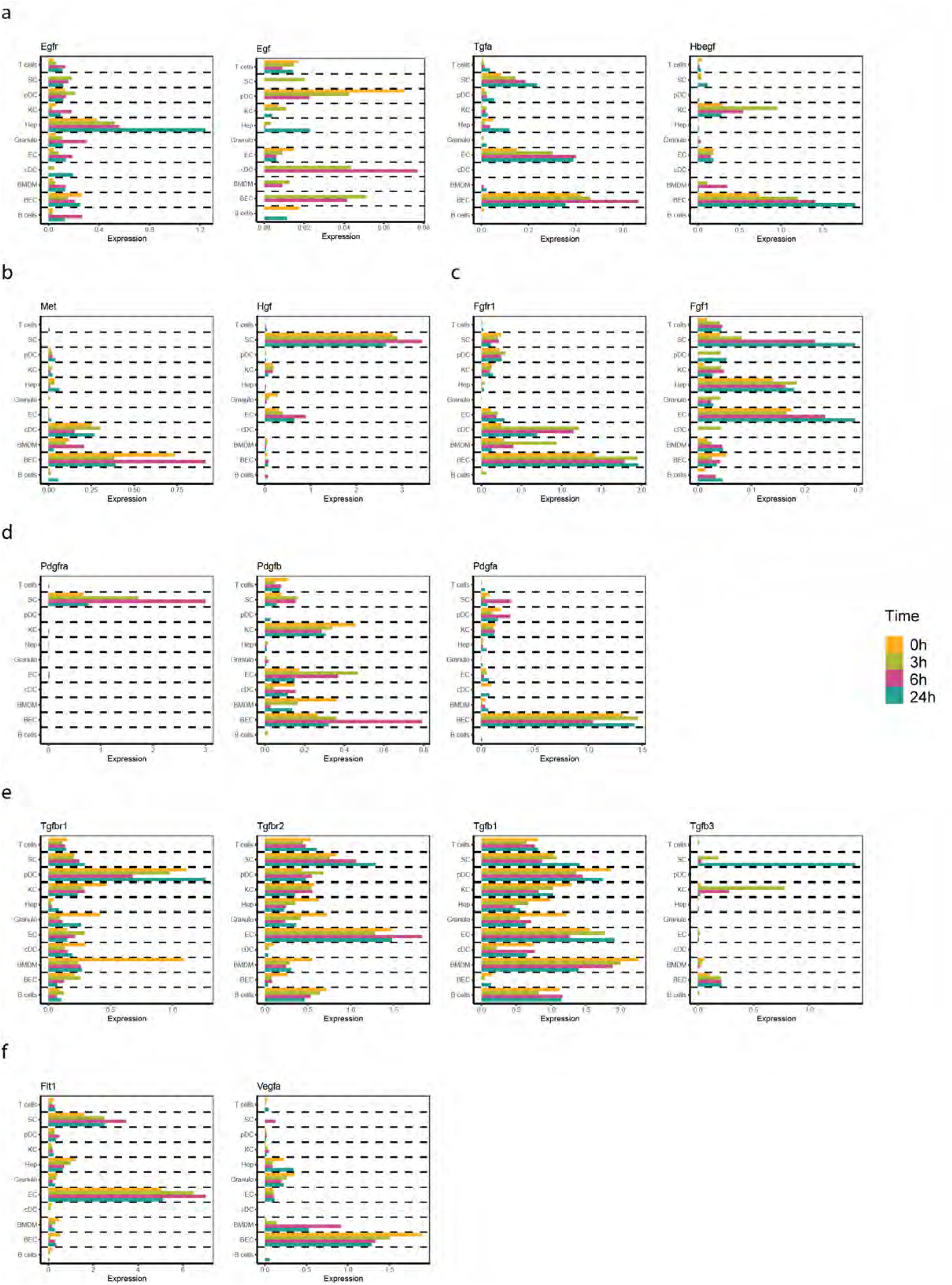
Gene expression of growth factors and growth factor receptors. Barplot of Egfr, Egf, Tgfa, Hbegf, Met, Hgf, Fgfr1, Fgf1, Pdgfra, Pdgfb, Pdgfa, Tgfbr1, Tgfbr2, Tgfb1, Tgfb3, Flt1 and Vegf.

**Extended Data Fig. 32.**
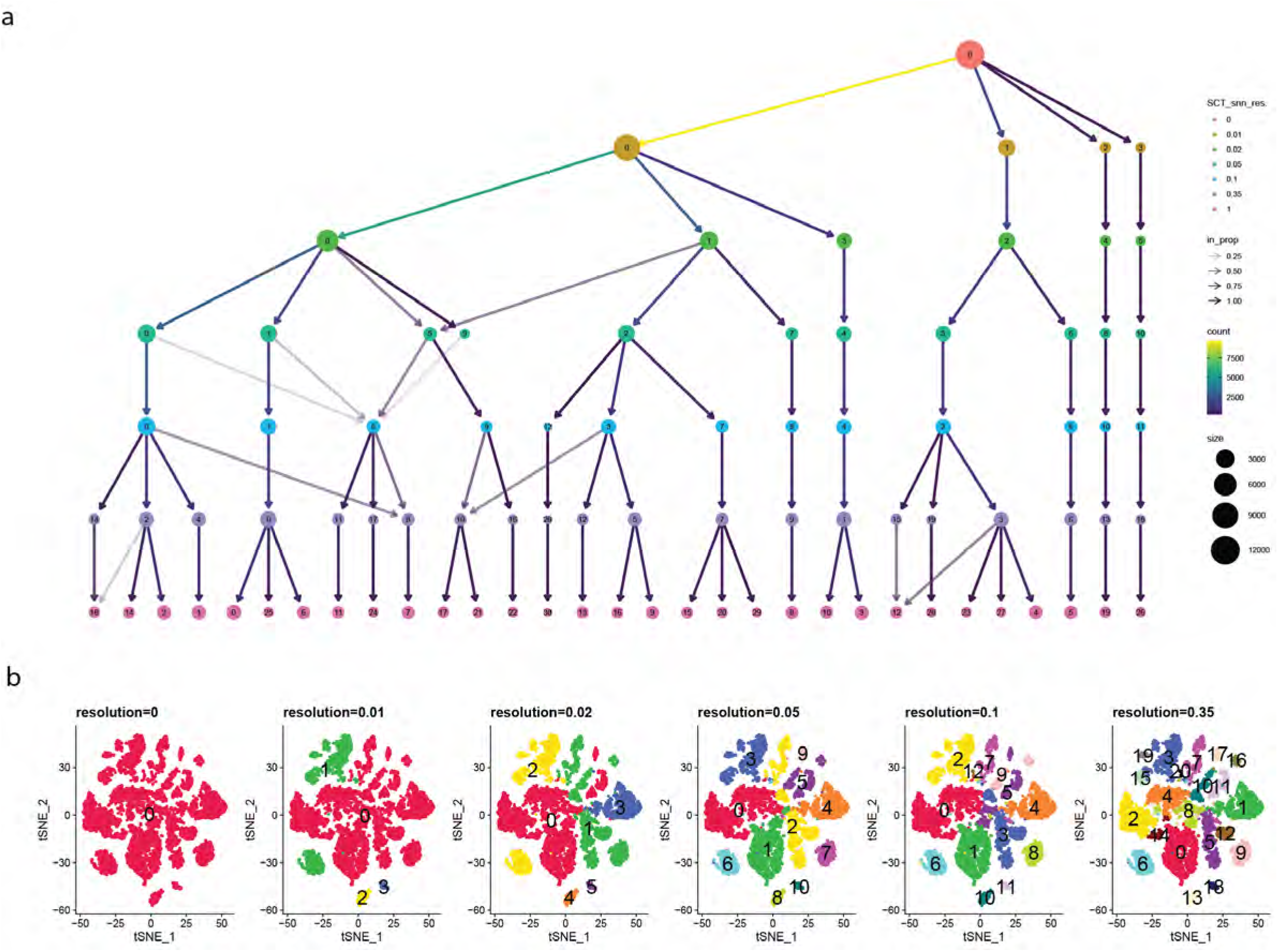
Emergence of clusters in scRNA-seq data. a) cluster tree with nodes colored by resolution, edges colored by number cells reassigned, node size represents the number of cells in a cluster and transparency is the proportion of cells reassigned. b) t-SNE visualization of the scRNA-seq data colored by unsupervised clusters with increasing resolution.

## Extended Data Table Legends

**Extended Data Table 1. Number of cells.** Number of cells per animal included for scRNA-seq and CyTOF after data preprocessing.

**Extended Data Table 2. Cell communication.** CellPhoneDB output showing the statistical significance of the interaction of the possible interactions between ligands and receptors including hepatocytes, endothelial cells, BMDM, Kupffer cells, B cells and granulocytes at 0, 3, 6 and 24 hours post PHx.

**Extended Data Table 3. Mitochondrial genes list and globin genes.** List of Mitochondrial genes used to identify dead cells in scRNA-seq (see methods) and list of globin genes used to identify red blood cells in scRNA-seq (see methods).

**Extended Data Table 4. Cell population marker. Markers used to identify each cell population in scRNA-seq data.**

**Extended Data Table 5. CyTOF panel.** CyTOF panel targets are organized by surface and cytoplasmic/nuclear staining compartments and in ascending order by metal conjugate.

**Extended Data Table 6. Flow Cytometry panel for identification of Kupffer cells and EdU incorporation.** Flow panel targets are shown in order of the staining protocol: first, surface targets, then intranuclear EdU staining and lastly the DNA intercalator.

